# A computational model of the spatiotemporal adaptation of tumor cells metabolism in a growing spheroid

**DOI:** 10.1101/2023.09.11.557115

**Authors:** Pierre Jacquet, Angélique Stéphanou

## Abstract

The Warburg effect, commonly depicted as an inherent metabolic trait of cancer in literature, is under intensive investigation to comprehend its origins. However, while the prolonged presence of excessive lactic acid production in tumors has been noted, it merely constitutes a fraction of the potential metabolic states cancer cells can adopt. This study aimed to elucidate the emergence of spatiotemporal diversity in tumor energy metabolism by expanding an existing model based on experimental facts. The resulting hybrid model integrates discrete formulations for individual cells and their processes, along with continuous elements for metabolism and the diffusion of crucial environmental substrates like oxygen, glucose, lactate, and the often underestimated acidity. This model enables simulation of a tumor spheroid, a standard experimental model, composed of numerous cells which can have distinct traits. By subjecting the spheroid to alterations of the environment such as cyclic hypoxia, acid shocks, or glucose deprivation, novel insights into metabolic regulation were obtained. The findings underscore the significance of the pyruvate-lactate interaction in governing tumor metabolic routes. Integrating acidity’s impact into the model, revealed its pivotal role in energy pathway regulation. Consequently, the conventional portrayal of a respiration/fermentation dichotomy proves inaccurate, as cells continuously and spatially adjust the ratio of these energy production modes, in contrast to abrupt, irreversible switches. Moreover, a cooperative cellular behavior akin to the reverse Warburg effect has emerged. This implies that the Warburg effect is not universally inherent to tumor metabolism, but a contextual, transient metabolic expression. Ultimately, the dynamic cellular-environment metabolic landscape influences cells’ survival under external conditions, with epigenetic regulations shaping their mobility potential within this landscape. While genetic mutations within tumor cells are undoubtedly present, this study shows they are not invariably essential for extreme metabolic modes or pathological characteristics to arise. Consequently, this research paves the way for innovative perspectives on metabolism, guiding tailored therapeutic strategies that consider not just patient-specific tissue attributes but also treat tumors as intricate ecosystems beyond their genetic diversity.

**Author Summary:** For years, scientists have been intrigued by the peculiar energy consumption patterns of cancer cells, such as the Warburg effect characterized by excessive lactic acid production. This study aimed to decipher the underlying reasons for the varying energy behaviors observed in different parts of tumors. Using a computational model, we simulated the collaborative dynamics of cells within tumors. The results revealed compelling insights. Two molecules, pyruvate and lactate, were identified as influential players in shaping energy utilization. Remarkably, the surrounding acidity was also found to exert a significant impact. Interestingly, tumor cells display a certain flexibility in their energy production strategies, adjusting according to prevailing conditions to maintain their survival and adaptability. Interestingly, cellular cooperation challenges the Warburg effect as an omnipresent phenomenon and reveals a transient nature. Our study underscores the significance of environmental influences, shedding light on the interplay between genetic modifications and the tumor environment in shaping cellular behavior. These findings hold promise for transforming cancer comprehension and devising treatments that tailor to both patients and the distinctive characteristics of their tumors.

## 1 Introduction

This work focuses on the adaptability of the cell metabolism in the face of environmental disturbances to which a tumor may be exposed. One of the manifestations of tumor metabolism, the Warburg effect, described almost a century ago, is often presented in modern literature as a universal characteristic of tumor metabolism. However, more and more experiments show that this is not systematically the case at the cellular level. In some recent works, we highlighted the semantic deviation associated with the Warburg effect, as well as concepts that are often attached to it such as *metabolic switch* or *metabolic reprogramming* [1, 2]. Far from neglecting the important role of mutations in the emergence of tumors, they cannot however explain all the observations specific to metabolism and its intrinsic properties and are not necessary to explain most metabolic behaviors.

What emerges from the literature is the need to integrate all biological scales to be able to address the complexity of tumor behavior. These approaches are increasingly put forward as the instrumentation evolves - in particular with multiomics approaches - and as the computing power increases. It also remains important to combine tissue and cellular measurements to account for differences in the dynamics [3]. If it is thus possible to claim that tumor metabolism is one of the defined states of a unified metabolic network, then it should be possible to produce a model capable of generating these different states.

In this work, a model of tumor metabolism highlighting the spatiotemporal heterogeneity is developed. This model makes it possible to test different scenarios of environmental stress and to assess their consequences on the metabolic trajectory of individual cells as well as of the tissue as a whole. Moreover, by combining continuous and discrete approaches within a hybrid multiscale model, cell metabolism is spatially contextualized and complex behaviors emerge.

In this new model, the environment is made of four diffusive species: oxygen, glucose, lactate and protons. The tumor tissue dynamics is represented by means of an agent-based model, where each cells composing the tissue is an agent having its own characteristics including its metabolite concentrations, gene expression levels, cell cycle, volume, *etc.* All of which are influenced by the local concentration of the environmental species *via* the cells incoming and outgoing flows. We recently developed a reduced model for cell metabolism, putting forward the role of acidity in the regulation of glycolysis and highlighting the rules for cell metabolic adaptation [4]. Our results suggested that what appears as a *cancer metabolic phenotype* is not necessarily due to cell abnormalities, but can spontaneously emerge as a consequence of the over-acidic environment [2]. Based on this earlier work, we propose here a more comprehensive model of cell metabolism inspired by the model of Li and Wang (2020) [5].

We extended their model to integrate the role of lactate and acidity. This includes the reversibility of lactate flows between the cell and the environment, the role of acidity on the direction of these flows as well as the already mentioned impact of this acidity on glycolytic activity. All these additions are based on recent observations from the literature [6, 7]. This new metabolic model was then implemented in the agent-based model and make it possible to address more complex metabolic states - involving cooperation between cells - and transitions in space and time.

The simulations presented, confirm our preliminary results by showing that the Warburg effect is not ubiquitous and temporality plays an important role. The expression of this effect is not always correlated with the fermentation activity of the cell. On the contrary, the metabolic state of the tumor cell is the result of a subtle balance between the environmental conditions and the internal cell conditions defined by its history. This state is not binary and is continuously adjusted using the means available to the cell allowing it not to stray too far from its homeostasis. It emerges from this model that these means are essentially thermodynamic in nature, *via* the enzymatic activity.

## 2 Models

### 2.1 A hybrid multiscale framework

Tumor spheroids are *in vitro* models of reference for the study of solid tumor growth [8, 9] because unlike conventional 2D monolayer cultures, they allow to study tumor heterogeneity. The cell density increases as they grow and this generates hypoxia which makes spheroids closer to *in vivo* conditions. Therefore, to address the heterogeneity of metabolism, we chose to build a spatio-temporal model of a tumor mass composed of thousands of cells, very similar to *in vitro* spheroid experiments, where each cell has its own internal metabolic dynamics. The models makes it possible to follow this very large quantity of cells over time, to observe the evolution of their phenotype and epigenome, to look at their distribution within the tissue, their energy profile and the impact of environmental conditions.

#### Spatial scales

The model is structured using a hybrid multiscale framework, integrating three main spatial scales (Fig.1) :

- the *environmental scale*, where the spatiotemporal dynamics of the constituents of the extracellular medium are described by mean of reaction-diffusion equations. The constituents in our model are limited to oxygen, glucose, lactate, and proton ions (for pH evaluation).
- the *tumor tissue scale*, where the cell assembly is described using an agent-based model. Each agent represent a cell and a predefined set of rules allow the cells to move, divide, interact or die.
- the *cell scale*, where each individual cell possesses its own characteristics, dynamics and fate. Specifically a mathematical model based on a system of reaction kinetics equations (mostly Michaelis-Menten kinetics) that reads as input the local environmental data, is solved for each cell.

**Figure 1:**
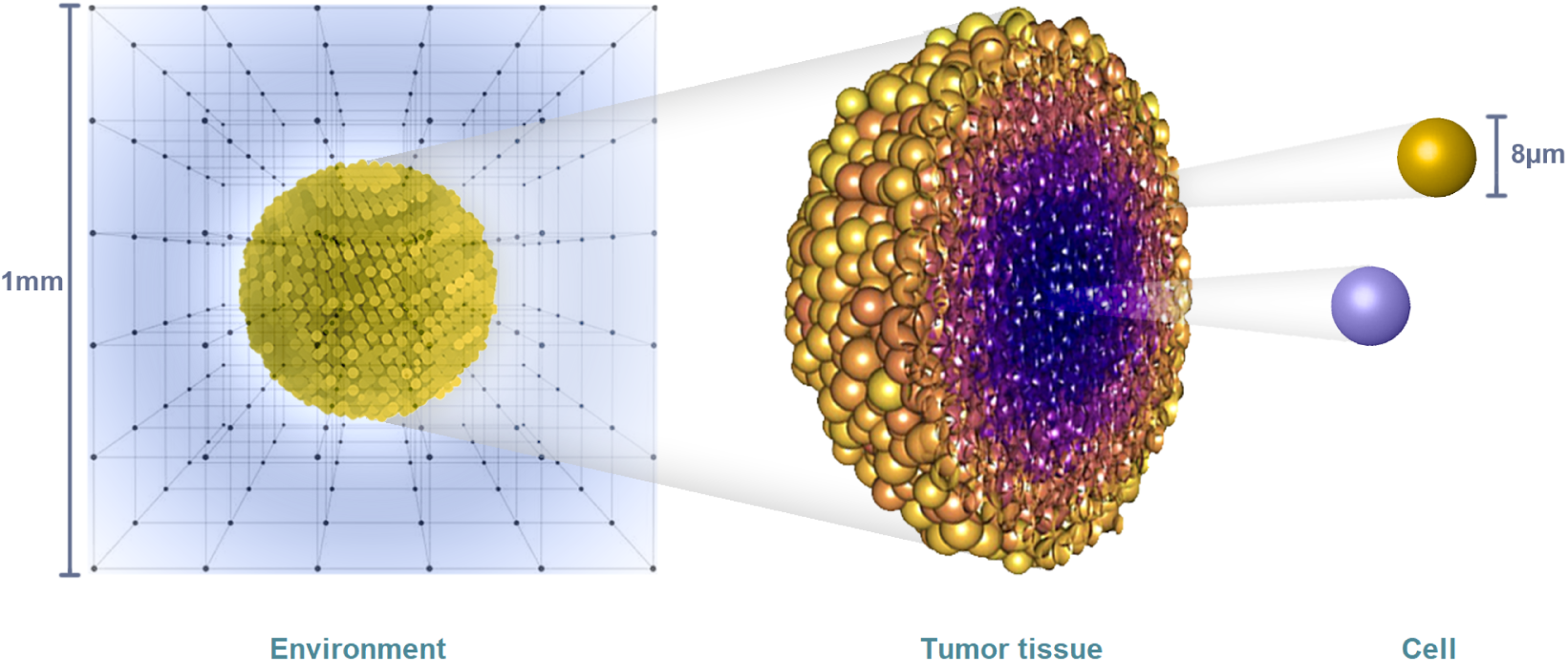
Structure of the hybrid multiscale model. The simulation domain is defined by a grid in which the PDEs for the four environmental variables are solved. The tumor tissue is made up of interacting agents, the cells, with free positioning (lattice free). Each cell has its own characteristics and dynamics.

We chose to use the *Physicell* software [10] for the implementation of our hybrid model. It allows to natively solve partial differential equations (PDEs) for the environmental variables, in a 2D or 3D simulation domain, using finite differences algorithms. The agent-based model (ABM) used to describe the cell behaviors integrates all the standard cell processes including the cell cycle, the cell division and cell mechanical interactions (adhesion or repulsion).

#### Temporal scales

The modeled processes do not occur on the same time scales, some processes being much faster than others. A different temporal resolution for each process makes it possible to modulate the relative impact of fluctuations occurring during very frequent processes, on less frequent processes. For example, the phenotype of each cell is evaluated only every 6 minutes of virtual time while their metabolism is evaluated every 1.5 seconds. In the *metabolism* frame of reference, the phenotype therefore appears to be stable. Another very important point is the saving in calculation time. If all the processes are evaluated at the smallest time scale, the computation time would be too long without obtaining more realistic results. Hence, the following time spans were used for each modeled biological process: 0.6 seconds for diffusion, 1.2 seconds for metabolism (resolution of differential equations = 2*×* diffusion), 6 seconds for mechanics (10*×* diffusion) and 6 minutes for phenotype (600*×* diffusion).

### 2.2 Modeling the extracellular environment

#### The substrates

In this study we focus on the cell energy metabolism. Therefore we chose to consider the three main following substrates to fuel the cell energy: oxygen, glucose and lactate. We moreover include protons as the fourth environmental variable of the model, since acidity plays a significant role in metabolic regulation that we recently highlighted in a reduced model of cell energy metabolism [4]. The range of values taken for each substrate variables are as follows:

- **Oxygen**: extracellular oxygen undergoes many variations within tumors. Oxygen is consumed first by the cells of the periphery of the spheroid leading to depletion at the heart of the tissue. The transport of oxygen inside the cells is done by passive diffusion, the intracellular concentration is therefore balanced with the extracellular concentration. The concentrations usually observed are 160 mmHg in the air at normal atmospheric pressure, 70 mmHg in the arteries, 38 mmHg on average in healthy tissues, the tissue is considered in hypoxia below 15 mmHg and in pathological hypoxia at 8 mmHg [11]. The initial concentrations chosen in the simulation - converted into mM using the Valabrègue coefficient 1.30 *×* 10*^−^*^3^ [12] - range between 0 and 38 mmHg. A constant oxygen value is prescribed at the domain boundaries (Dirichlet’s condition), since in experiments, spheroids are continuously exposed to air, unless otherwise specified. The oxygen diffusion coefficient is 87,600 *µ*m^2^*/*min [13].
- **Glucose**: like oxygen, glucose is depleted by the cells. Its transport inside the cells is by facilitated diffusion involving a GLUT transporter. The blood glucose concentration is close to 5-6 mM in average. A constant concentration of glucose is prescribed at the domain boundaries (Dirichlet’s condition), since the culture medium is regularly renewed in spheroid experiments, unless otherwise specified. The glucose diffusion coefficient is 30, 000*µ*m^2^*/*min [14].
- **Lactate**: lactate is mainly produced by the cells and consumed to a lesser extent. Lactate enters and leaves the cells via MCTs (monocarboxylate transporters) in the form of lactic acid. For one lactate molecule leaving the cell, one H+ ion also leaves. At the domain boundaries, the concentration of lactate is most of the time fixed at zero, since the extracellular medium is regularly renewed in spheroid experiments. In some of the simulations the constant boundary condition (Dirichlet’s condition) is replaced by a zero flux condition (Neumann’s condition) allowing the lactate produced to accumulate. The lactate diffusion coefficient is 12,600 *µ*m^2^*/*min [15].
- **H+**: protons have the same dynamics as lactate. They enter and exit cells in the model, only via MCTs. The resulting pH often ranges from physiological (*∼* 7.4) to very acidic (*∼* 4.0). The pH at the domain boundaries is fixed at 7.4 and like lactate can be replaced in some simulations by zero flux conditions. The protons diffusion coefficient is 270, 000*µ*m^2^*/*min [16].

#### Non-homogeneous diffusion

In the *Physicell* version that we used (version 1.7.1), the solvers for diffusion equations were only considering a constant diffusion coefficient. The initial simulations that we did, were not satisfactory as we could not generate realistic oxygen and nutrients gradients in the spheroid. We therefore modified the solver to integrate the case of non-homogeneous diffusion. Specifically, we introduce a dependency with regards to the local cell density ρ to better reflects the gradients intensity. The non-homogeneous diffusion is expressed as:

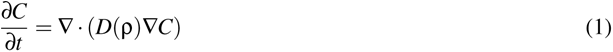

where *C* is the diffusive species. For each voxel, the cell density is evaluated (in % cell occupancy of the sampled volume). If the local density is 1 (100% of the surrounding voxel volume is occupied) then:

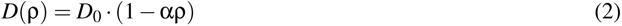

with *D*_0_ the diffusion coefficient of the species in the interstitial liquid. α *∈* [0, 1] is a parameter for adjusting the impact of the density on the diffusion, and ρ is the local density of the cells in the voxel. By using α = 0.25, the diffusion profiles obtained for the three substrates and pH (Fig.2) resemble those found in the literature [17].

**Figure 2:**
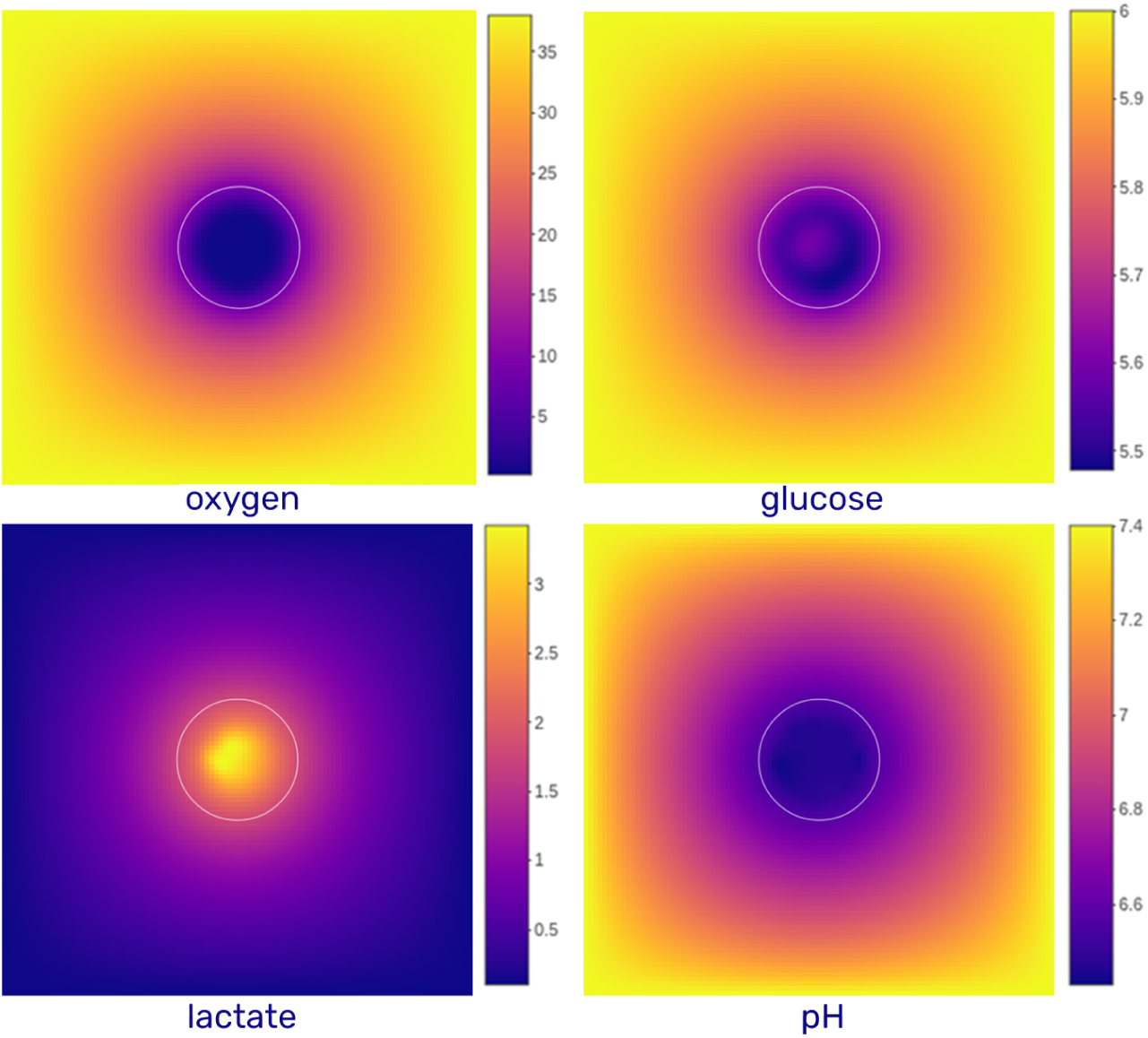
Diffusion profile for the environmental variables. The diffusion domain is a grid of 2000 *µm* of side (divided into 100 elements of 20*µm*). The white circles materialize the contour of the spheroid. The color gradient represents the concentration of each substrate. Oxygen is expressed in mmHg, glucose and lactate are expressed in mM and pH is without units (pH is calculated from the proton concentration expressed in mM). Constant values are maintained at the domain boundaries for all variables (Dirichlet’s conditions).

### 2.3 Modeling the cell metabolism

We recently proposed a reduced model for cell metabolism [4] that highlights the role of acidity as a regulator of the cell metabolism. The model we propose here - although more complex - retains the main hypotheses that we put forward in the reduced model. Those are detailed below.

#### 2.3.1 The model by Li and Wang

The mathematical model that we propose is derived from the model by Li and Wang, 2020 [5]. In this article, the authors extended and refined the model of Marín-Hernández *et al.*, 2011 [18] which focused on glycolysis, by including the mitochondrial energy pathways, the concentrations of different enzymes, as well as the regulation of ten key genes.

The objective of this model was to look at the interaction between the gene regulatory network expressed with ODEs and the metabolic expression. This model has more variables and is finer in the description of the reactions than our previous reduced model [4], but it keeps the same overall modeling philosophy. The parameters are based on experimental data and the system as a whole is numerically stable. The system of differential equations integrates fluctuations and small random disturbances inherent in real biological systems. The equations become stochastic and instead of evaluating the evolution of the trajectory of the different variables (concentrations of oxygen, glucose, expression of the p53 gene, *etc*.), it is the evolution of their distribution that is considered. The *landscape* of stationary states and their probabilities is then maped [5].

Due to the large number of variables which results from this, all the dimensions of the system do not need to be treated at the same time because the authors have chosen to look only at the landscape formed by two variables which are the concentrations of PDH (Pyruvate dehydrogenase) and LDH (lactate dehydrogenase). These two enzymes are somehow representative of the bifurcation between the two main metabolic modes (fermentation and respiration).

However, in its current version, the model was not fit to address some of our questions regarding spatiotemporal metabolic heterogeneity. In particular, the time evolution of the different cellular configurations was not considered. Instead, the probability of states *via* the topology of the attractors was privileged. Thus it was the derivative at time *t* of each flow for a large number of conditions which was evaluated. This only gave a static image of the landscape disconnected from the context of tumor growth. Without temporal tracking, the extracellular values do not need to be dynamically updated. The pH which shapes the landscape over time was not integrated. Finally, the spatial distribution of metabolism was not addressed.

As a consequence, we took over the equations of this model by adding different aspects that were not covered by the authors, specifically, the temporality, spatiality, adjustment of lactate transport mechanisms and influence of the pH.

#### 2.3.2 Model adaptations

The biochemical model of metabolism takes up and modifies that of Li and Wang [5]. The original model is written in MATLAB code. It has been translated into symbolic code in Python, which is used to directly generate the C++ code of the system of equations as well as the associated Jacobian matrix, for direct integration into *PhysiCell*.

The differential equations describe the one- or two-way reaction flows and take the following form [19] which is an extended form of a Michaelis-Menten equation, reversible and extensible to several substrates [*S*] or several products [*P*]:

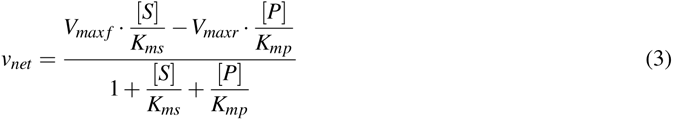

here *V_max f_* and *V_maxr_*are respectively the maximum speeds of the reaction in the direct (forward) and reverse (backward) directions. *K_ms_* and *K_mp_* are the Michaelis constants of the substrate and the product respectively.

Each cell within the simulation integrates the entire system of equations corresponding to its metabolism with its own parameters and concentrations. At the start of the simulation, the few initial cells have slightly different internal concentrations for each metabolite (around the values in the literature). At each time step (*dt_ODE_* = 1.2sec), the set of cells is taken, in a random order and for each, its ODE system is solved with the Rosenbrock solver of order 4, from the C++ Boost Odeint library [20] that was integrated into *PhysiCell*.

All the equations are available in Supporting Information (S1 Appendix). They are for the most part very close to those proposed by Li and Wang [5]. Table 1 recapitulates the metabolic reactions of the model and highlight those that were modified.

For most of the fermentation reactions (in green in table 1), the expressions have been multiplied by a term *pH_inhibition_*which decreases the activity of the enzymes when the pH becomes acidic. The enzymes chosen (and their associated reaction) are HK, GPI, PFK-1, ALD, TPI, GAPDH, ENO, LDH, G6PD-6PGD, PFKB2/3. These correspond to the enzymes whose activity was affected by the pH in the study conducted by Xie *et al.* [6]. Based on their observations the term *pH_inhibition_* has been added:

**Table 1:** Metabolic reactions integrated into the model. Green reactions are reactions that have undergone minor changes compared to the basic model, in orange major modifications.

| IDs | Enzymes | Reactions |
| --- | --- | --- |
| r1 | GluT1 | $\text{Gluout} \rightleftharpoons \text{Gluin}$ |
| r2 | HK | $\text{Gluin} + \text{ATP} \rightleftharpoons \text{G6P} + \text{ADP}$ |
| r3 | GPI | $\text{G6P} \rightleftharpoons \text{F6P}$ |
| r4 | PFK-1 | $\text{F6P} + \text{ATP} \rightleftharpoons \text{FBP} + \text{ADP}$ |
| r5 | ALD | $\text{FBP} \rightleftharpoons \text{DHAP} + \text{G3P}$ |
| r6 | TPI | $\text{DHAP} \rightleftharpoons \text{G3P}$ |
| r7 | GAPDH | $\text{G3P} + \text{NAD}^+ \rightleftharpoons \text{1,3BPG} + \text{NADH}$ |
| r8 | PGK | $\text{1,3BPG} + \text{ADP} \rightleftharpoons \text{3PG} + \text{ATP}$ |
| r9 | PGAM | $\text{3PG} \rightleftharpoons \text{2PG}$ |
| r10 | ENO | $\text{2PG} \rightleftharpoons \text{PEP}$ |
| r11 | PKM2 | $\text{PEP} + \text{ADP} \rightleftharpoons \text{Pyruvate} + \text{ATP}$ |
| r12 | LDH | $\text{Pyruvate} + \text{NADH} \rightleftharpoons \text{Lactate} + \text{NAD}^+$ |
| r13 | G6PD | $\text{6PGD G6P} \rightleftharpoons \text{R5P}$ |
| r14 | ATPases | $\text{ATP} \longrightarrow \text{ADP}$ |
| r15 | AK | $\text{AMP} + \text{ATP} \rightleftharpoons 2\text{ADP}$ |
| r16 | PFKFB2 | $3 \text{ F6P} \rightleftharpoons \text{F2,6BP}$ |
| r17 | PHGDH | $\text{3PG} \longrightarrow \text{Serine}$ |
| r18 | PDH | $\text{Pyruvate} + \text{ADP} \longrightarrow \text{Citrate} + \text{ATP} + \text{Complex2}$ |
| r19 | ACC | $\text{Complex2} + 3\text{ATP} + \text{AC-CoA} \longrightarrow \text{mal-CoA} + 3\text{ADP} + \text{NAD}^+$ |
| r20 | SOD | $\text{ROS} \longrightarrow \text{Null}$ |
| r21 | | $\text{Lactate} + \text{H}^+ \rightleftharpoons \text{Lactate}_{extra} + \text{H}^+$ |
| r22 | | $3\text{R5P} \rightleftharpoons 2\text{F6P} + \text{G3P}$ |
| r23 | NUCLEOTIDE<br>BIOSYNTHESIS | $\text{R5P} \longrightarrow \text{Null}$ |
| r24 | SERINE<br>CONSUMP-TION | Serine → Null |
| r25 | GPDH | NADH + ADP → Complex2 + ATP + NAD+ |
| r26 | | Citrate + 3ADP → 3ATP + 4Complex2 |
| r27 | | Complex2 + 1.5ADP → 1.5ATP |
| r28 | | Complex2 → ROS |
| r29 | NOX | null → ROS |
| r30 | | Citrate → Null |

**Table 2:** General trends of simulations as a function of external conditions. . The oxygen, glucose and lactate concentrations are reported qualitatively and the results column indicates the consequences of these conditions on the exposed cells.

| Oxygen | Glucose | Lactate | Results |
| --- | --- | --- | --- |
| + | + | ++ | Lactate import |
| + | + | + | Lactate excretion |
| + | + | - | Low lactate excretion |
| + | - | + | Lactate import |
| + | - | - | Death |
| - | + | + | Lactate excretion |
| - | - | + | Death |
| - | - | - | Death |

#### Inhibition term by pH

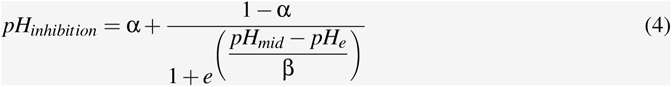

with α, the minimum level of enzymatic activity at acidic pH (set from [6] at 0.1= 10%), β defines the regulation sensitivity (set at 0.12) and *pH_mid_* is the pH at which the enzyme operates at 50% of its maximum activity (fixed at 6.7).

The addition of this term makes it possible to integrate recent observations of a reduction in the Warburg effect when this has led to the acidification of the environment. This simple addition is a source of heterogeneity within the tumor, particularly with respect to the gradients of exposure to acidity.

In the original model, oxygen is considered as a parameter with a fixed value and has no direct impact on OXPHOS. The lack of oxygen increases the production of HIF, which triggers a series of hypoxic responses, but the respiratory chain functions normally, as long as enough complex II units of the chain have been formed. However, in reality oxygen is required as the final electron acceptor. This is why the two ATP-generating expressions (r18 and r25), involving complex II (in the initial model), are multiplied by a Michaelis-Menten term which strongly reduces the reactions below a hypoxic threshold [11] (0.011 mM *O*_2_).

#### Term for inhibition by oxygen

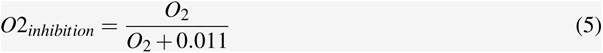

Since we are here in a situation where the oxygen evolves significantly (as it is consumed by the cells) and we are interested in the trajectory of the system over time, oxygen is no longer a simple parameter but a variable. Thus, it was necessary to calculate the quantity of oxygen consumed from the activity of the respiratory chain, that is to say how much ATP it produces. Modeling mitochondrial metabolism is also a complex exercise and requires a very large number of equations if one wishes to integrate each mechanism involved.

The model of Li and Wang [5] integrates the essential mechanisms to model the flows that interest us at the cost of a division of reactions that is not necessarily very intuitive: to produce ATP in the mitochondria, *NADH* and *FADH*_2_ are each oxidized but do not produce the same amount of protons in the intermembrane space. The *NADH* produces a little more which allows *in fine* to make the ATPase work and produce a little more ATP (approximately 2.5 ATP for *NADH* and 1.5 ATP for *FADH*_2_). In the model the *FADH*_2_ is not taken into account, the authors therefore establish, in a first reaction (r25), the account of the excess ATP produced by *NADH* (in relation to *FADH*_2_, *i.e.*, one molecule of ATP) thus creating the “complex II” mitochondrial unit. And, in a second step, they calculate (r27) the remaining quantity of ATP (= 1.5) commonly produced by *NADH* and *FADH*_2_. This division avoids having to integrate an additional variable and simplifies the number of equations to be solved, although this makes the equations less easy to read.

It is currently estimated that on average (again it depends on many conditions) for 12.5 ATP generated in the mitochondria from a molecule of pyruvate, 80% comes from *NADH* (10 ATP) and 20% of *FADH*_2_ (1.5 ATP). If we reduce this calculation to the number of complex II formed, we therefore have: 1.5 *ATP_FADH_*_2_ *·* 0.2 + 2.5 *ATP_NADH_ ·*0.8 = 2.3 *ATP/complex II* in the electron transport chain.

Moreover, the quantity of oxygen consumed during the transition from complex II to complex IV also depends on the proportion of *NADH* and *FADH*_2_ used. With *NADH*, the ATP/oxygen ratio (P/O ratio: ATP formed per oxygen molecule consumed) is 5, against 3 for *FADH*_2_. So we have 1 ATP for 1/5 *O*_2_ with *NADH* and 1 ATP for 1/3 *O*_2_ with *FADH*_2_. Keeping the same proportion as before *NADH/FADH*_2_, we therefore have: 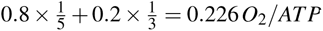.

The final formula for the oxygen consumed can therefore be summarized as follows:

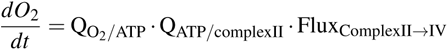

That is to say :

#### Oxygen over time

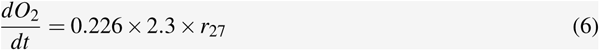

#### Flows entering and leaving the cell

Lactate transport between the cell and the environment has been revised to incorporate proton transport. In the initial model, this depends only on the intra/extra-cellular lactate concentration gradient. However, monocarboxylate transporters (MCT1 & MCT4 here) work closely with proton dynamics [21]. The transport therefore does not only depend on the lactate gradient but also on the protons, so that the final dynamic results from a sort of equilibrium between the two.

The transport equation was composed based on the following experimental observations made by Xie *et al.* [6] :

- whatever the pH, the ratio of intracellular lactate/pyruvate concentrations is greater than approximately 10;
- the pyruvate concentration is kept relatively constant and remains above 0.1 mM;
- when the extracellular pH is acidic and the extracellular lactate concentration is higher than the intracellular one or the lactate/pyruvate ratio falls too low, lactate is transported into the cell;
- when the extracellular pH is neutral and even if the extracellular lactate concentration is higher than the intracellular one, very little lactate enters the cell.

Biologically, MCT1 and MCT4 are distinguished with respect to their thermodynamics, one and the other allowing easier transport in one of the two directions (as shown in figure 3 with the case of lactate). To simplify the process and avoid having to quantify the number of carriers of each type, only one generic carrier responding to the global dynamics is modeled:

**Figure 3:**
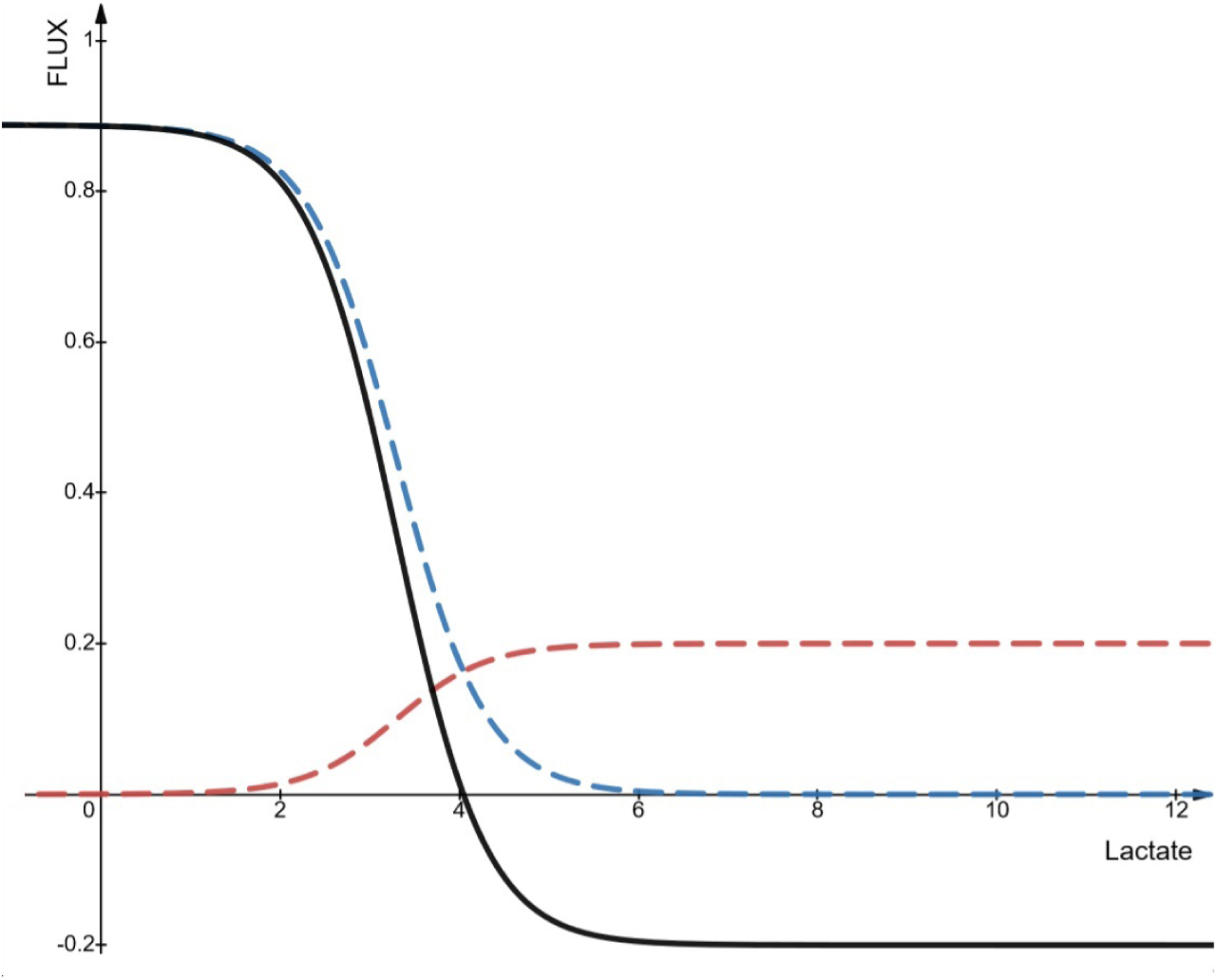
Incoming and outgoing flux of lactate and protons according to the extracellular concentration of lactate. In blue, the outgoing flux , in red the incoming flux (absolute values) and in black the resulting flux (r21). In this figure the concentration of pyruvate is 0.11 mM, intracellular lactate at 3.3 mM, the extracellular pH at 6 and the intracellular pH at 6.7 and *V_m f_*_21_ =1

#### Lactate and protons flux

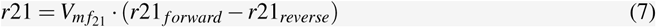

#### Lactate and protons flux

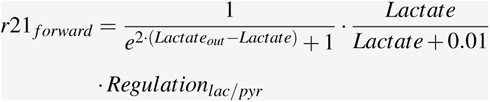

with *Lactate_out_* the concentration of extracellular lactate and *Lactate* the intracellular one.

The first term of *r*21 *_f orward_*, makes it possible to reduce the outgoing flow when the extracellular lactate concentration is close to or higher than the intracellular one and, conversely, to bring it to 1 when *Lactate > Lactate_out_* . The second term cancels the first when the intracellular lactate is close to zero (so as not to artificially release lactate which does not exist).

The last term manages the dynamic between lactate and pyruvate and results in:

#### Regulation of the ratio between lactate and pyruvate

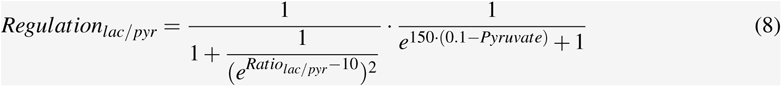

with

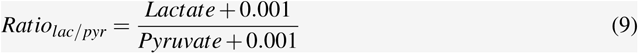

The lactate/pyruvate ratio (*Ratio_lac/pyr_*) is calculated by adding very small terms of 0.001 (compared to standard concentrations) so as not to have critical values, if the pyruvate concentration becomes zero. The first term of *Regulation_lac/pyr_* makes it possible to vary the outgoing flow of intracellular lactate from 1 (when the ratio is greater than 10) to 0 when the ratio drops below. The second term makes it possible to increase the outgoing flow of intracellular lactate from 0 to 1, when the concentration of pyruvate is greater than 0.1 mM, in order to keep the lactate inside the cell if the pyruvate becomes rare. The value of 150 is an arbitrary value to adjust the steepness of the curve so that the flow increases quickly. However, this value cannot be too high so as not to create transitions that are too abrupt and difficult to solve for numerical solvers.

It should be noted in these last equations the presence of coefficients which have not been put in the form of parameters to facilitate reading because of the large number of parameters already present in the system. Moreover, these values are considered constant, once they have been fixed (like 150 for example). Also, the equations can be written in a more compact form, we chose not to do it, to keep track of the modeling choices and to facilitate their explanation. In the code, these equations are simplified automatically using the symbolic calculation software Sympy [22].

#### Incoming transport of lactate and protons

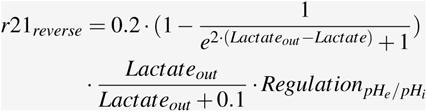

The first term of the incoming transport is a constant fixed here at 0.2 and which is linked to the maximum speed of the incoming flows by the MCT transporters. If the maximum outflow value is 1, the maximum inflow is only 20% as efficient in the experiments of Xie *et al.* [6]. Again this parameter could be adjusted but it was not necessary here to do it dynamically. The second term is the counterpart of the first term in the outflow equation. It is subtracted from 1 in order to increase the inflow when the concentration of extracellular lactate is close to or greater than the intracellular one and conversely to bring it to 0 when *Lactate > Lactate_out_* . The third term cancels the second when the extracellular lactate approaches 0 so as not to transport lactate inside when it is not present. The last term *Regulation_pH__e/pHi_* is defined by:

#### Regulation of the inward transport of lactate and protons by pH

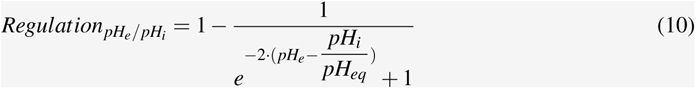

This last term varying between 0 and 1, promotes (the term approaches 1) the incoming transport of lactate and protons when the difference between *pH_e_*and *pH_i_*weighted by *pH_eq_* is positive. Typically the value of *pH_eq_*is 1.1 which allows an equilibrium for an extracellular pH slightly more acidic than the intracellular one.

Figure 3, shows the result of modeling r21 for *V_m f_*_21_ = 1. It can be noted for example that the final flow in black is slightly below its maximum speed of 1 when the extracellular lactate is close to 0 because the pyruvate is very close to its minimum concentration and the lactate/pyruvate ratio is not large enough to compensate. Export is therefore important due to a lack of lactate on the outside, but not maximum to avoid depleting all the pyruvate. The pH being sufficiently acidic, the inflow (in red) tends towards its maximum speed of 0.2.

To calculate the concentrations of the different metabolites, all that remains is to add the corresponding incoming and outgoing fluxes (table 1). For example for intracellular glucose:

#### Concentration of intracellular glucose over time

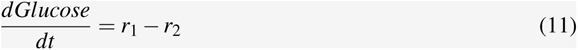

For protons, the only fluxes having an impact are those of lactate transport (r21). As specified above, for an excreted or integrated lactate molecule, a proton accompanies it. We would therefore tend to think that:

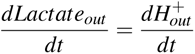

In reality, this is not really the case because of the phenomenon of buffering (buffer effect of a solution). When a small amount of proton is added to a solution, it is able to withstand a change in pH so close to an equilibrium between weak acids and their conjugate bases. The buffer capacity is a measure of this resistance and is specific to each type of solution. Van Slyke’s equation [23] makes it possible to correctly calculate the evolution of the pH of a solution according to the existing pH and the quantity of protons added (via r21). The following form follows directly from the application of Van Slyke’s equation:

#### Evolution of the concentration of extracellular H+ ions over time

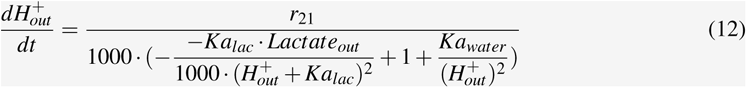

with *Kal_ac_* = 10^−3.86^ mole/L, dissociation constant of lactate and *Ka_water_* = 10^−14^ mole/L, dissociation constant of water. The constants of 1000 are here to convert molars into millimolars.

A note on the implementation in *PhysiCell*: when several cells are in the same place (above the same voxel), they read the same concentrations at a time *t* and attempt to transport the same fluxes. In the event that there are not enough nutrients for all the cells in the voxel, the risk is that the concentrations become locally negative, *i.e.* the cells have ingested more nutrients than available at a moment ***t***. To avoid this, a “voxel debt” system has been integrated, which instead of transferring the matter between the cells and the voxel all at once, spreads this quantity until the next time step at which the metabolism is evaluated. The simulations generated are no different from the simulations for which a smaller time step would have been taken.

#### Regulation of gene expression

In the previous section, the equations defining metabolic fluxes were defined. Some of these equations are directly influenced by the expression level of different genes (for example the glucose transporter GLUT1 in the r1 reaction). They can also be influenced indirectly by other genes which act upstream (for example HIF increases the production of GLUT1 which in turn acts on r1). The model by Li and Wang [5] thus integrates 13 genes acting closely with the enzymes and metabolites of the model. To study the dynamics of their expression, their level is normalized.

#### Evolution of the expression level of a gene over time

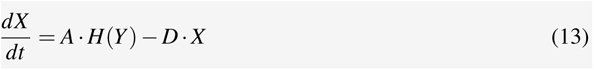

The expression level of each gene *X* increases at a basal rate A of 0.005/min (normalized level/min) and is degraded at the same rate *D*. These values are calibrated on measurements carried out in previous studies and specified in the related publications. *H*(*Y*) is a Hill function defining the regulation of the gene *Y* on the gene *X*:

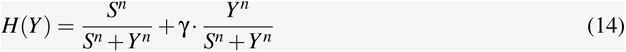

γ defines the type of regulation (positive > 1, negative < 1 and neutral =1), *S* the gene level *Y* at which the expression is half (similar to *K_m_* of a Michaelis Menten) and *n* the regulatory force (between 0.025 and 40).

All these equations are also defined in the Appendix S1. No changes have been made to these equations apart for that of HIF. It is degraded at a constant rate in the original publication. We made it depend on the oxygen level by adding the term *H*(*O*_2_) to the HIF degradation term.

What primarily defines the behavior and regulation of the genes is the regulatory strength parameter γ. The closer the γ of a gene-gene, gene-enzyme or metabolite-gene pair is to a normalized value of 1, the greater the effect of the first member of this pair on the other member will be relatively weak, and conversely the further away it is, the stronger it will be. The value of γ was generated randomly in the original model. Since we are considering the trajectory of the metabolism over time, we chose to introduce the transmission of a close value of this parameter to the daughter cells after cell division.

Thus, the cells at the start of the simulation are initiated with values of γ close to each other and calibrated from the literature. When a cell divides, for each γ of each gene in its system, new random values are drawn following a normal distribution centered on the current values of γ (by preserving the mode of action, that is to say that if the gene has an inhibitory action the random value will not be drawn above 1). The new values are therefore more likely to change little, but major upheavals can occasionally occur. This allows to carry out a progressive drift of the epigenome of each cell. If the new configuration is not viable, the cell will eventually die and the propagation of these new γ will be limited.

### 2.4 Modeling the cell cycle and cell death

#### Cell cycle

Each cell in the simulation has a cycle, evaluated at each *dt_phenotype_* that determines what it does or cannot do. The cell cycle is calibrated on U87 human glioblastoma cells which have an approximate cycle time between 18 and 24 hours [24]. The first G1 phase has a basal duration of 5 hours. When the cell is in this phase it evaluates (checkpoint 1 of Fig.4) if the level of ATP is sufficient to continue its cycle. The cell needs energy to prepare and initiate division, if it does not have enough, continuing the cycle is risky. In addition, it assesses whether the cell density around it is not too high. If one of the two criteria is not met, it enters the G0 phase (quiescence phase) and remains there for an indefinite time until the conditions are favorable again (checkpoint 2). Unlike healthy cells, tumor cells divide even when there is little room to do so. However, this phenomenon has limits and although they are higher than for healthy cells, tumor cells cannot endure infinite density. A reference density has therefore been fixed and the cells undergoing a higher density enter the G0 phase.

**Figure 4:**
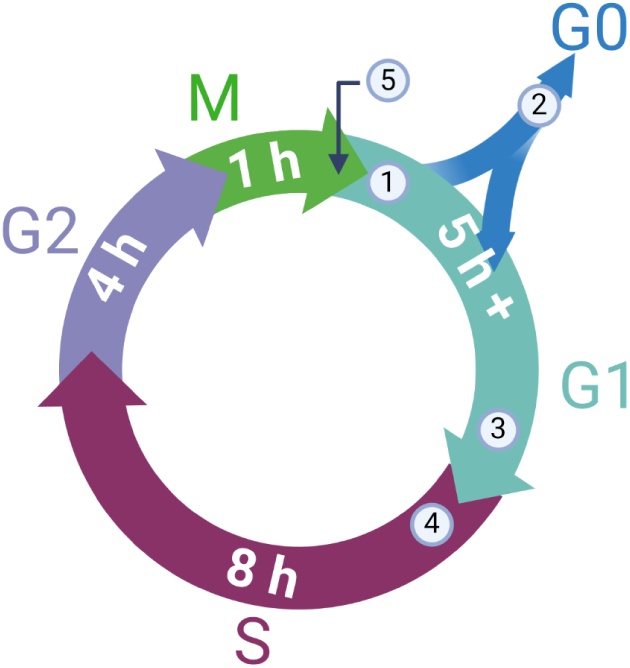
Cell cycle used in the simulation. Control points are indicated by circled numbers. 1) Evaluation of the transition to G0. 2) Evaluation of return to G1. 3) Latency of transition to S phase. 4) Doubling of cell volume. 5) Cell division.

During the G1 phase, if the oxygen level drops (without reaching hypoxia), the phase will lengthen by a few hours in proportion to the oxygen level, in order to delay the irreversible transition to phase S in case conditions deteriorate (checkpoint 3) [10]. When the cell enters the S phase it begins to double its current volume (checkpoint 4) in preparation for future division. At the end of the G2 phase, when the cell enters the M phase, the values of the γ parameter described in the previous section are chosen. This corresponds to epimutations which makes it possible to introduce internal variability gradually into the cells. Finally, at the end of the M phase, the cell having doubled in volume splits into two daughter cells of identical volume, each having the same parameters and concentrations but which will now operate independently.

All the cells therefore have a common cycle, but on the scale of the entire population, this cycle is asynchronous. The cells are not all in the same phase at the same time. Thus at the start of the simulation, when the cells are initiated, the phase of the cycle for each of them is drawn randomly with a weight proportional to the duration of each phase.

#### Cell death

Necrosis and apoptosis are the two types of death a cell can undergo. Tumor cells have acquired the ability to escape apoptotic death but are not unconditionally immortal. In this model, the parameter that determines cell death is ATP. If the cell happens to be below a critical ATP threshold, it has a high probability of inducing necrotic death. The more the level of ATP is below the threshold, the more this probability increases. This threshold is arbitrarily set at 0.3 mM (the conditions of death by ATP depletion vary widely - some studies mention between 15 and 25% [25] of the basal level of ATP, which approximately brings the threshold to this value). To protect the cell from death caused by a very momentary loss of ATP (environmental hazards or nutrients taken at time *t* by one or more neighboring cells), death is only possible if this deficit is maintained over time (3 hours in the model).

When the cell dies by necrosis, it enters a first stage during which its volume increases (*necrotic swelling*), then in a second stage the volume decreases again until the cell disappears in the form of debris (the cell is destroyed and is removed from the simulation). During this last phase, glucose, oxygen, lactate and intracellular protons are released into the medium where they diffuse freely.

## 3 Results and discussion

The simulations performed aimed at establishing first a context of reference that represents a typical *in vitro* experiment of the growth of a spheroid. The simulation was fully characterized to provide a reference state from which to compare simulations made in different contexts. Perturbation of the environmental conditions were then considered including cyclic hypoxia, acid shock and glucose deprivation. Finally, we challenged the glycolytic metabolism to verify if and how the cancer cells are able to adapt to new conditions. The results are presented in the next three dedicated sections.

### 3.1 Reference simulation

Spheroids represent a good *in vitro* model for tumor growth [26] since (*i*) spatial heterogeneities due the limited diffusion of the nutrients and oxygen can be established from the periphery to the core of the spheroids; (*ii*) it is easy to manipulate the environmental conditions such as oxygen or glucose concentrations. Therefore, we chose to simulate the case of a spheroid grown in a round bottom low attachment well as shown in figure 5.

**Figure 5:**
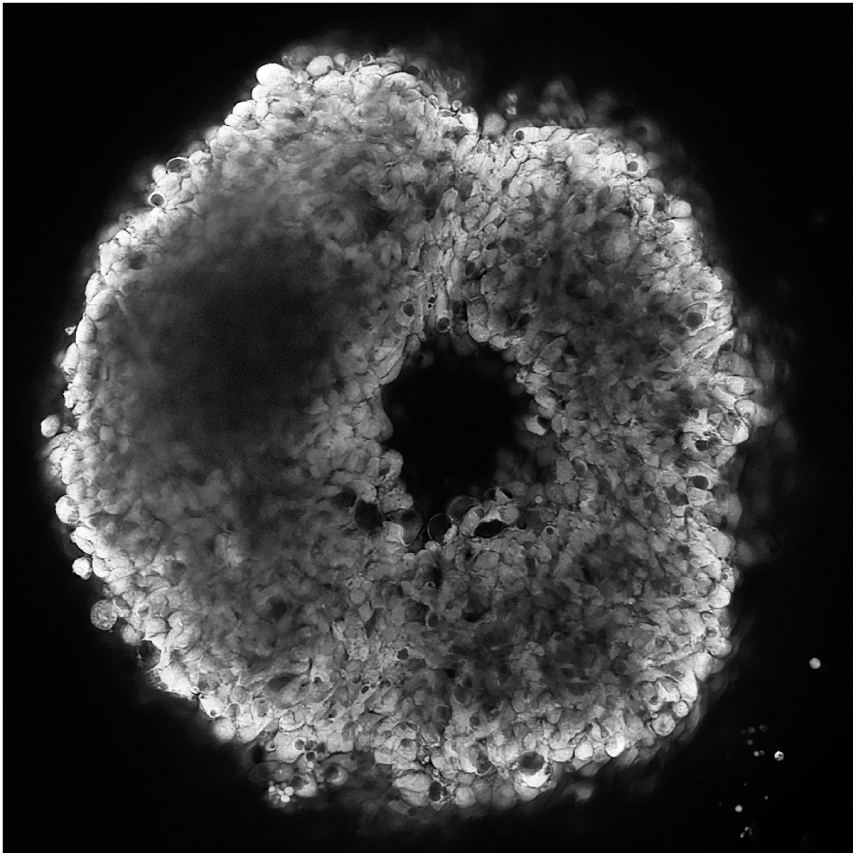
Spheroid of F98 cells. This spheroid, approximately 390 *µm* in diameter, was obtained from F98 cells (rat glioma). Cells are marked with the fluorescent probe BCECF, which makes it possible to measure the intracellular pH. The center of the spheroid which exhibits no fluorescence is a necrotic core (dead cells).

#### 3.1.1 Initial conditions

To establish the reference simulation, the glucose concentration at the start of the simulation is set at 6 mM. Oxygen is set at 38.0 mmHg which corresponds to the pressure measured in *in vivo* tissues to better capture the reality of the metabolism. The lactate is in very small quantity with 0.1 mM and the pH is fixed at the physiological value 7.4. For the three substrates (oxygen, glucose and lactate) and the pH, the values taken as initial conditions are kept constant at the boundaries of the simulation domain (Dirichlet’s condition) as if the culture medium was continuously renewed, to maintain glucose concentration and eliminate lactate and as if the pH level was maintained with buffer solutions. Oxygen is also kept constant passively by contact with the environmental gas conditions set in the incubator.

Although the simulations can be made in 3D, we will only consider 2D simulations because of the high computational cost. As a consequence, the cells can only move on the plane and the “2D spheroid” diameter will expand faster than the real 3D one. The goal here is not to quantitatively compare the spheroids size and growth rate but simply to find common emerging structures to appreciate their similarities.

Figure 6 presents the reference simulation with the initial and boundary conditions previously defined. At time *t* = 0*h* a small spheroid of 440 cells was considered. At the end of the simulation (ended as the cells come into contact with the domain boundaries) at time *t* = 443*h* the final number of cells was about 50000. The spheroid morphology remains homogeneous as it grows. A necrotic core progressively develops and the dead cells are progressively eliminated from the simulation domain in *Physicell*, leaving a hole very similar to the one observed experimentally (Fig. 5).

**Figure 6:**
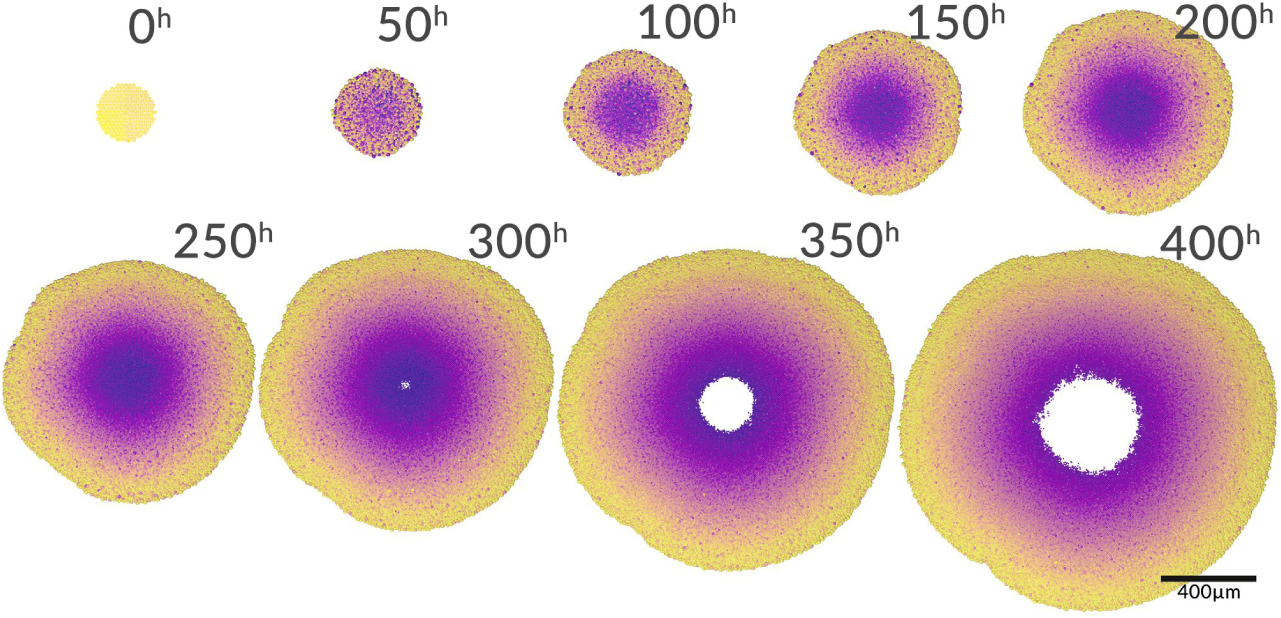
Reference simulation of spheroid growth. Each image is spaced 50*h* from *t* = 0 to 400*h*. The color code is relative to each time step and indicates the age of the cells. The more purple a cell is, the older it is relative to other cells.

#### 3.1.2 Gradients and radial profiles

The main advantage of a spheroid is that it is a simple tissue model with radial symmetry. Thus the majority of the emergent heterogeneity is along this radial profile. This comes from the exposure to different gradients of substrates that are differentially processed (produced or consumed) from the periphery to the core of the spheroid.

The first emerging profile is that of the distribution of cellular states (Fig.7). In this figure and on the spheroid growth graph (Fig.8), the spheroid begins its growth with only proliferating cells. None entered the G0 phase and the external conditions allow the development of the tumour. At 50*h*, the appearance of an isolated cell layer, of quiescent cells, is observed. This comes from the stabilization of cell density. The cells no longer have room to grow and the infusion of nutrients begins to decrease. At 100*h*, a necrotic core appears. The conditions of oxygen, glucose and pH, although relatively favorable outside the spheroid, are not sufficient to compensate for the lack of oxygen or the glucose depletion as well as the acidity which settles in the center. This necrotic core persists until the end of the simulation. From 300*h*, the most central cells died and turned into debris, leaving a hole instead.

**Figure 7:**
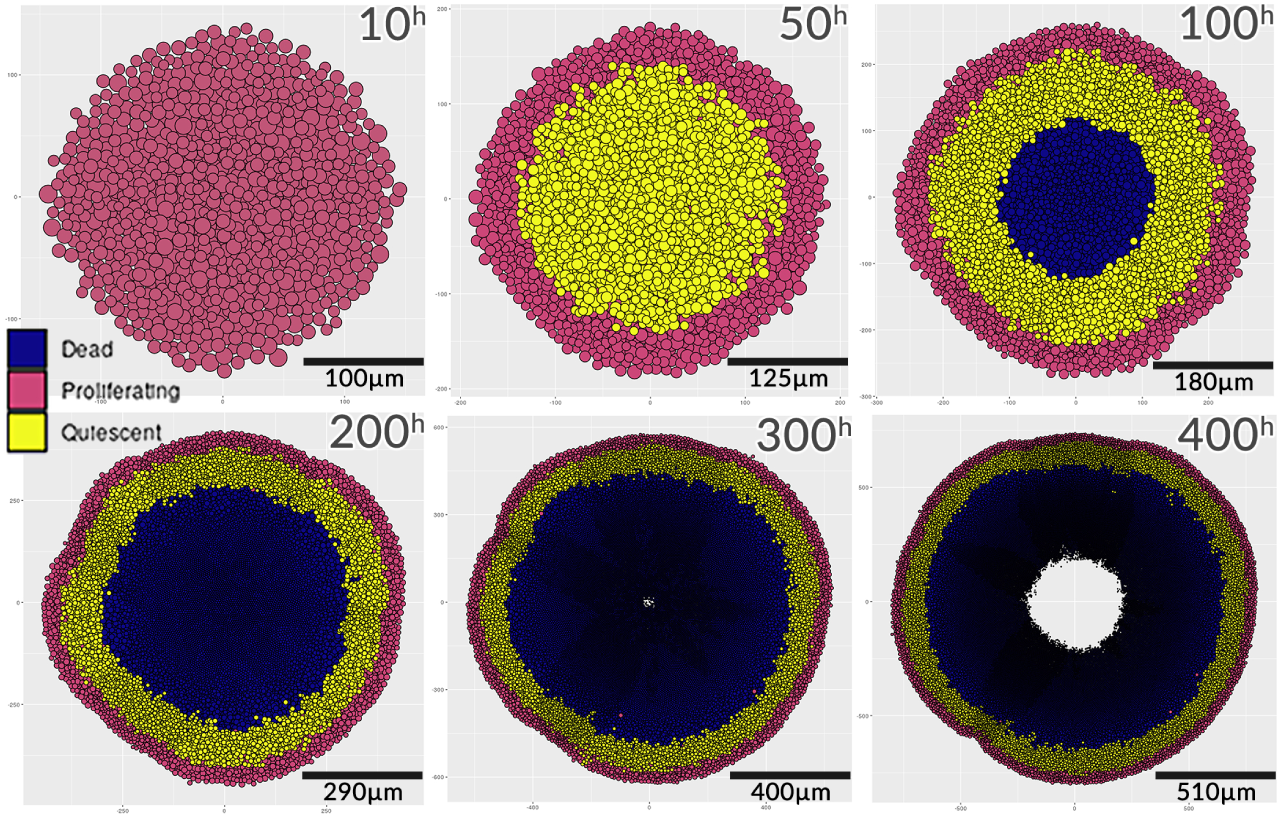
Distribution of the cell states of the spheroid in the reference simulation over time. The six figures present the evolution of the simulation from 10 to 400*h*. In blue the cells have started the necrotic process, in yellow the cells are in a state of quiescence (G0) and in pink, the cells are in a proliferative state (G1,S,G2,M). In black, cellular debris are gradually removed from the simulation.

**Figure 8:**
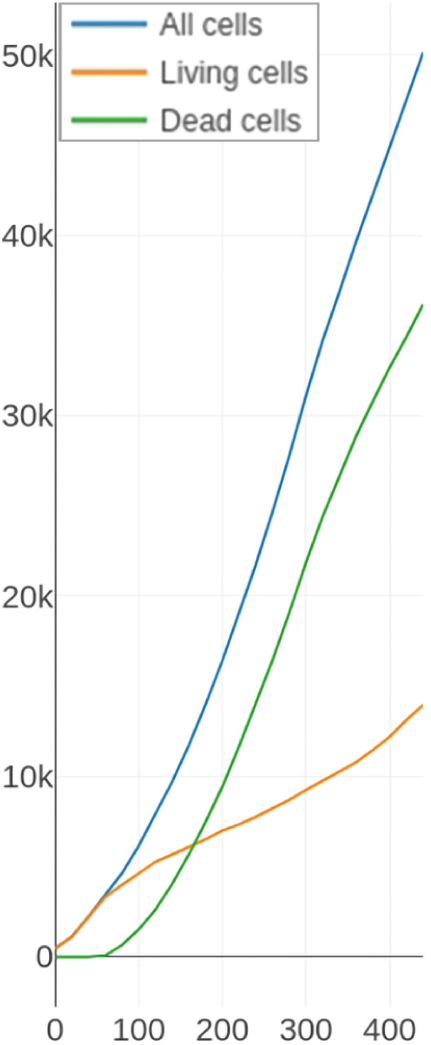
Evolution of the different cell populations in the spheroid for the reference simulation.

Figure 9 shows the evolution of the concentrations of the substrates in the medium surrounding the spheroid in the reference simulation. For each substrate, the value of the concentrations present in each voxel is averaged with those of the other voxels having the same distance from the center of the spheroid. These concentration profiles as a function of the distance to the center are calculated for each time step of the simulation.

**Figure 9:**
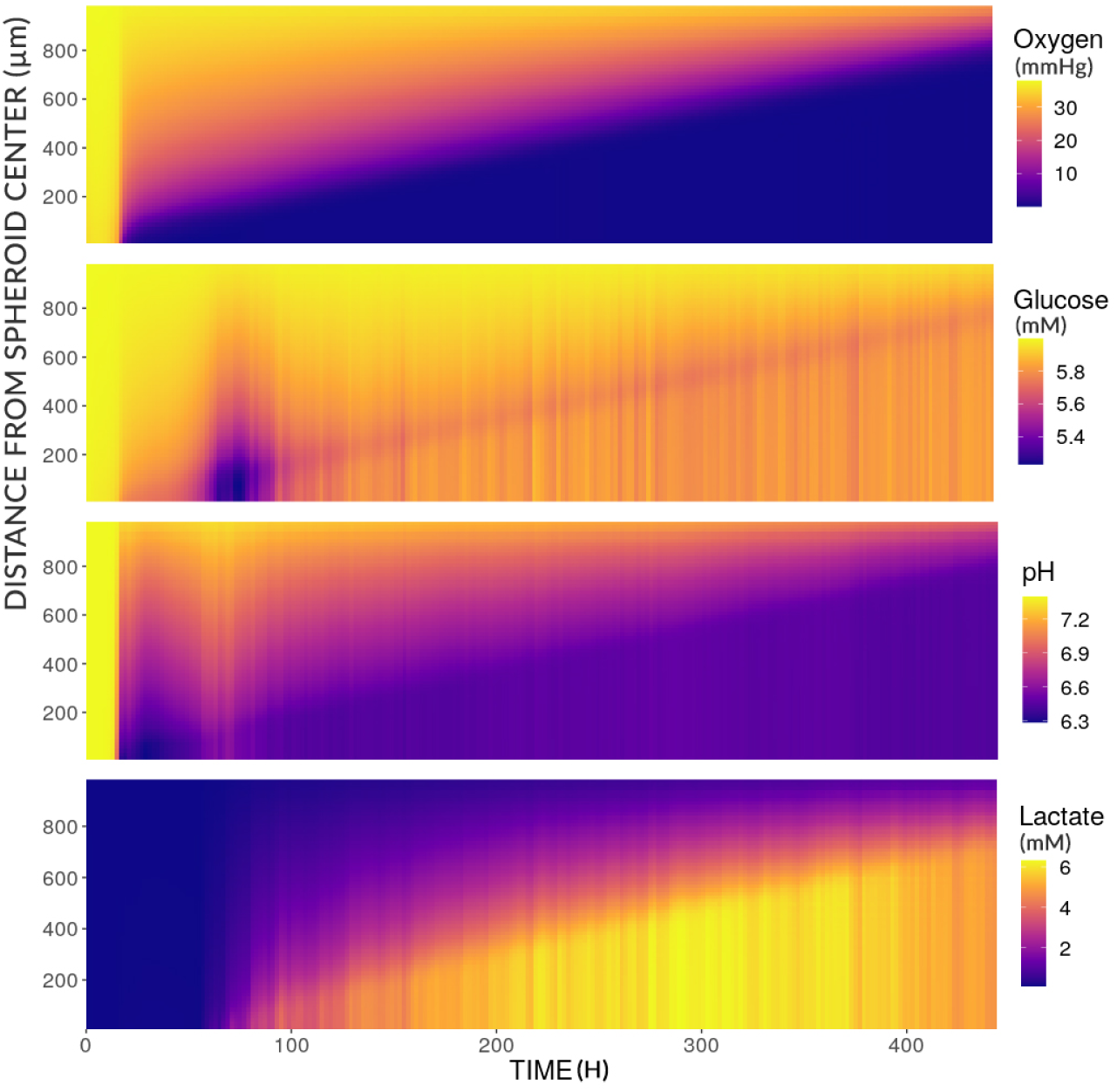
Mean radial profiles of substrate concentrations in the environment of the reference simulation over time.

The figure thus shows that up to approximately 20*h* of simulations, the diffusion rates of oxygen and glucose are sufficiently rapid and the consumption of the cells too low to cause visible depletion in the medium (yellow area). This time roughly corresponds to the time necessary for all the cells in the simulation to have completed at least one complete cell cycle (and therefore a division leading to the increase in density). Oxygen depletion then increases in a quasi-linear manner, leaving mainly only the first layers of cells to have full access to oxygen. The inner layers can also have access to oxygen but in less quantity, systematically depleting all the molecules that reach them. The pH also decreases indicating glucose consumption and lactate production.

Between 50 and 100*h*, the pH very briefly becomes neutral again and the glucose concentration suddenly decreases. This indicates a lack of pyruvate and a compensation for this lack by recovering the little lactate in the medium (and therefore protons). By raising the extracellular pH, glycolysis is no longer inhibited and glucose consumption increases again. However, lactate is not present in large quantities and only glycolysis then makes it possible to maintain the necessary levels of pyruvate that can be used by the mitochondria. This re-causes a drop in pH in the inner layers of the spheroid again inhibiting glycolysis.

From 100*h*, lactate is present in greater concentration, but it can no longer be used to replenish pyruvate within the cell, since the latter is no longer consumed anyway due to lack of oxygen. As a result, only the layers on the periphery of the spheroid have access to oxygen and although glucose is available, it is not enough to produce the level of ATP necessary to keep the cells in the deepest layers alive. Cells in the peripheral layers use oxygen and glucose (shown by the purple line in the glucose profile), producing lactate and acidity in the inner layers.

#### 3.1.3 Emerging phenotypes

Figure 10 presents the metabolic phenotypes of the reference simulation. To assess the cell phenotypes, fluxes involving pyruvate, at the intersection between glycolysis, lactate and acetyl-CoA, are measured. Here what is observed is the distribution of pyruvate fluxes in respiration or fermentation and not the resulting amount of ATP. Thus, if the flux value of r12 (reaction catalyzed by lactate dehydrogenase - see table 1) is greater than the flux value r18 (reaction catalyzed by pyruvate dehydrogenase) then the phenotype is essentially associated to fermentation, with a little respiration (in light red) on the figures. If r12 is 10*×* greater than r18, *i.e.* more than 90% of the pyruvate is directed to lactate production, then the phenotype is pronounced fermentation (dark red). The opposite situation r18 greater than r12 (and 10*×* greater) gives the respiration phenotype (purple) and pronounced respiration phenotype (dark purple).

**Figure 10:**
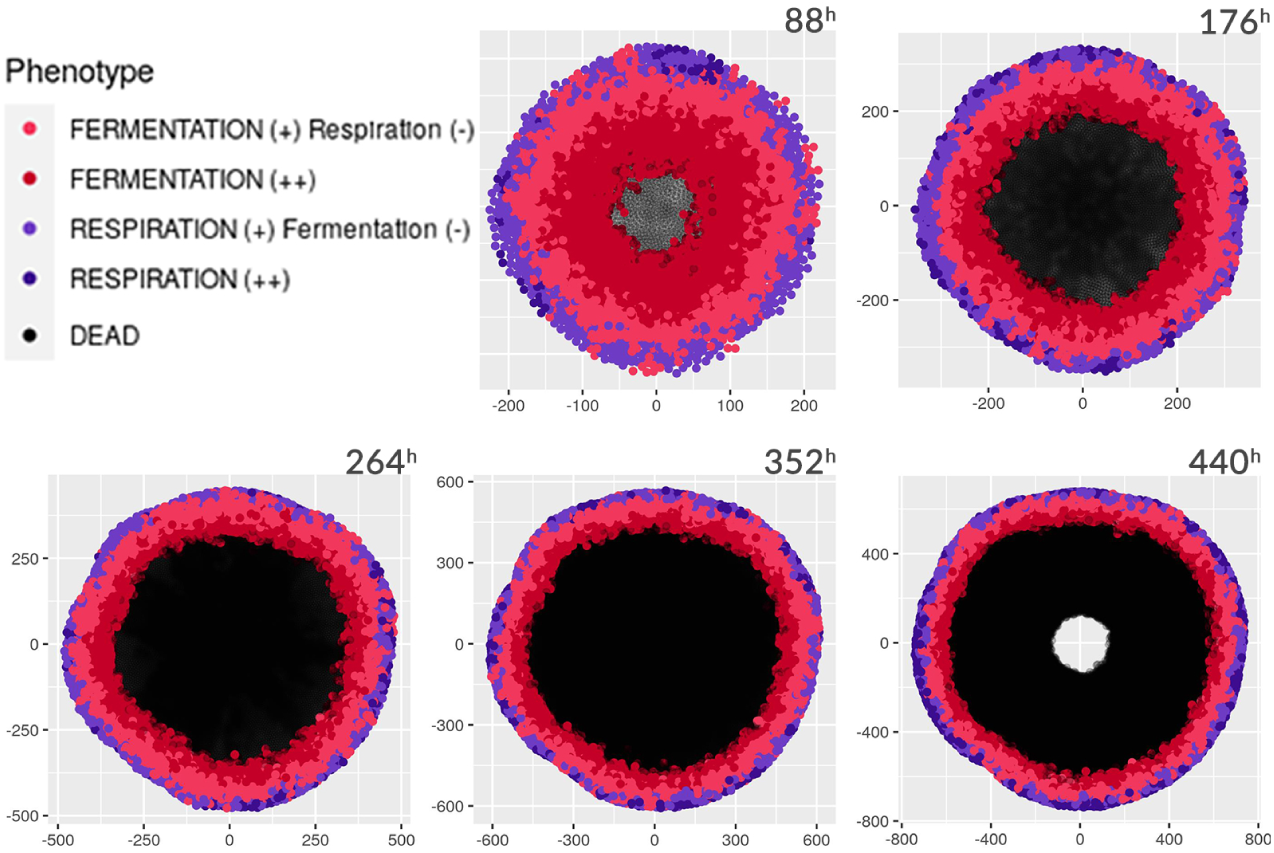
Spheroid cell phenotypes from the reference simulation over time. The cells are represented here with an identical size in order to better visualize the phenotypes. When a process is described with a (-) it means that it is present but in a lower proportion than that marked with a (+). The black cells are in necrosis.

Throughout the simulation, the outermost layer of cells retains a predominantly oxidative metabolism often coupled with a little fermentation (purple cells). Some cells (dark purple) have a pronounced oxidative metabolism but it does not last long. The cells in the layers below begin to ferment more than they respire, the lack of oxygen as well as the lactate coming from the lower layers favoring this state. Finally, the cells still alive in the deepest layers ferment exclusively when there is no oxygen available.

Looking now at the lactate fluxes and in particular by labeling these phenotypes by the manifestation of the Warburg or reverse Warburg effect, the results may seem surprising.

In figure 11, two definitions of the Warburg effect are represented [2]. The modern definition (in light orange), corresponding to the production of lactate in the presence of oxygen (thus indicating that pyruvate is transformed into lactate rather than into acetyl-CoA). The original definition corresponding to the simple excretion of lactate in the tumor is represented in dark orange. We note that no minimum threshold is defined to qualify an excretion of lactate as corresponding to the Warburg effect. Finally, the reverse Warburg effect (concretely the import of lactate by the cell) is in light blue.

**Figure 11:**
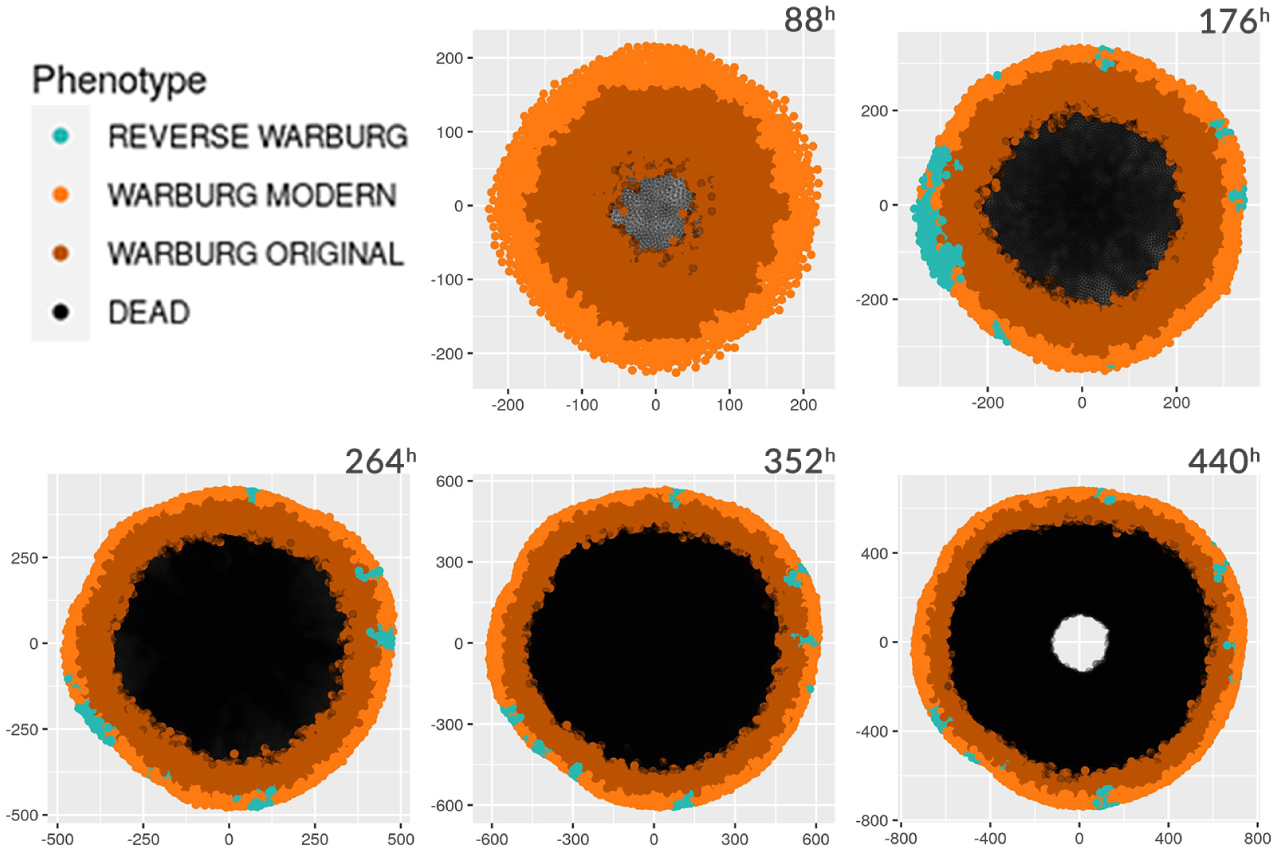
Warburg and reverse Warburg effect in the reference simulation of spheroid cells over time. The black cells are in necrosis.

The Warburg effect appears to dominate tumor cell phenotypes with few exceptions of the reverse Warburg effect. This indicates that even cells previously labeled with *respiration* or *pronounced respiration* excrete lactate. In fact, the latter secrete it in much smaller quantities than cells in hypoxia. In reality, it is not because lactate is produced by the cell that it is excreted and the reverse also. Experiments carried out on the relationship between intracellular and extracellular lactic acid concentration [6] show that overall these concentrations are balanced. A cell having thus accumulated lactate at a given moment can excrete it later when the gradient is reversed while at the same time having a respiratory metabolism. The question therefore arises of the criteria necessary to precisely qualify the Warburg or reverse Warburg effect, when observed at the cellular level and on the relevance of such criteria.

#### 3.1.4 Metabolic Landscape

In Li and Wang [5] from which the model presented is derived, metabolic attraction zones are described but also transition phases between these zones. Metabolic states are described as basins continuously linked to each other. Four pools of attractions are identified (*normal, oxidative cancer, glycolytic cancer, intermediate cancer*) by the expression levels of LDH and PDH genes. However, we here chose to replace *normal* by *neutral* so as not to force a distinction between a healthy and cancerous mode. By choosing these two genes, it is the bifurcation at the level of the fate of the pyruvate which is evaluated. The interest for looking at the levels of these genes in relation to the reactions on which they act (respectively r12 and r18), is that the variation in their level of expression is “slower”. This makes it possible to look at the medium term effects of the environment on the metabolism of the cell, rather than its immediate response to environmental fluctuations.

It is therefore interesting to look at the changes in the topology of this landscape, but mainly by looking at the capacity of the cells to move around, that is to say to reach the identified zones.

Figure 12 shows that the cells move within three of the four zones described. At the start of the simulation, the cells move towards two fairly well-defined zones: the glycolytic zone at the bottom right of the landscape (strongly expressed LDH) and towards the oxidative zone at the top left (PDH more expressed). The distribution observed previously in figure 10, between fermentation and respiration phenotypes is very clear here (particularly at 20*h*). The acidification of the medium coupled with the oxygen still present at 30*h*, leads to the transition of a large part of the cell population into a momentarily more oxidative mode for about twenty hours. Figure 13 highlights this transition.

**Figure 12:**
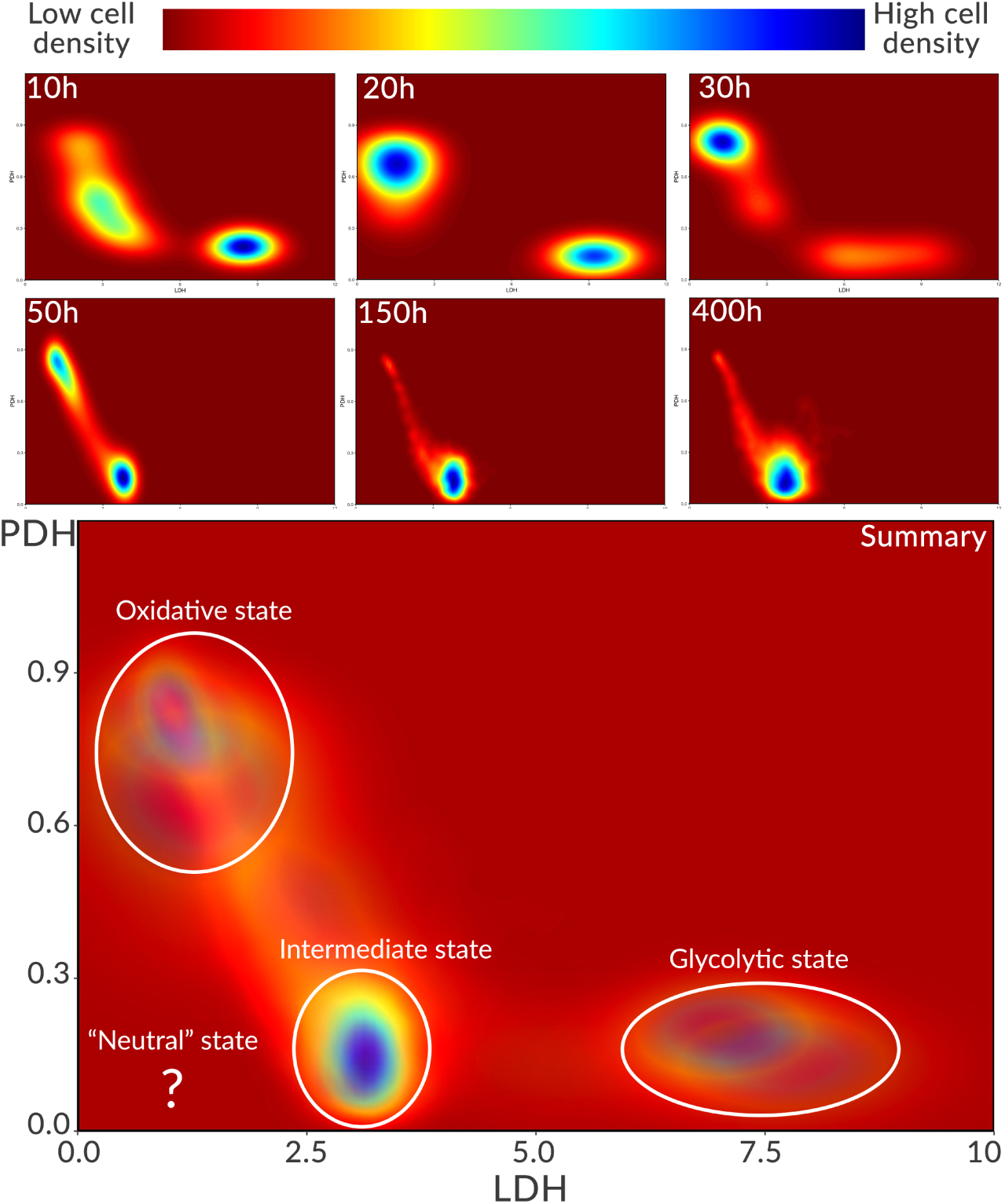
Evolution of the metabolic landscape of the tissue of the reference simulation over time. The maps present the metabolic density zones (number of cells), at the level of the expression of the LDH and PDH genes. The red areas are areas where very few cells are found, unlike the blue areas which contain a large quantity. The map below is the aggregation of the metabolic landscape over time, accumulating areas of high densities. The white circles delimit these areas. The *neutral* zone is the only zone that does not appear on this map.

**Figure 13:**
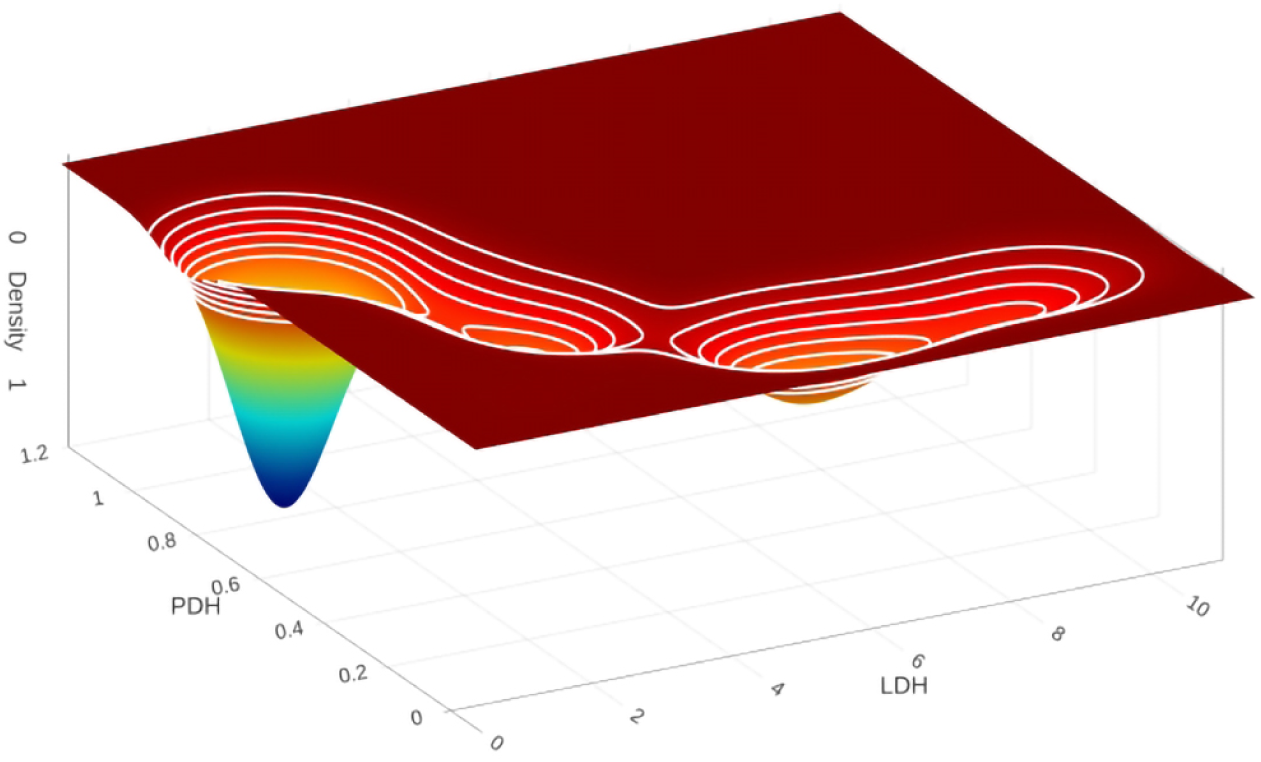
Distribution of cell density in the LDH/PDH bifurcation from the reference simulation to. 30*h*. The white lines, represent lines of iso-density within the metabolic landscape.

However, the growth of the tissue quickly leads to a significant depletion of oxygen, which no longer allows the expression of PDH to be maintained as high. From 50*h*, the cells stuck between the lack of oxygen and the increasing acidity are pushed into a hybrid state between the *intermediate* state and the *neutral* state. This general tendency is maintained until the end of the simulation and the transition between the *oxidative* phenotype of the cells in the periphery and the *intermediate* state is clearly visible.

### 3.2 Environmental perturbations

#### 3.2.1 Cyclic hypoxia

Cancer cells are often exposed to cycles of hypoxia, notably caused by a complex and constantly reconfiguring vascular network [27]. Experiments to show the effect of these variations have been conducted with tissues intermittently exposed to oxygen tensions ranging from 10 mmHg to 60 mmHg (some up to 160 mmHg but these values are not physiological) [28]. These oscillations can have very fast (less than an hour), intermediate (tens of hours) or slow (greater than a day) frequencies [28, 27]. It is interesting to look at the repercussions of such oxygen oscillations on metabolism and to look at the ability of cells to adapt more or less quickly to these changes. For this, three simulations were carried out based on the conditions presented in the reference simulation. However, here the oxygen at the edges of the environment is successively fixed at a high concentration (70 mmHg) then suddenly falling at a low concentration (15 mmHg) corresponding to hypoxia. These values are slightly higher than the values cited above to take into account the depletion gradient settling in the spheroid. These successions of normoxic and hypoxic states are carried out cyclically at a frequency of 18, 36 and 72 hours (each state then lasting 9, 18 and 36 hours). The other parameters of the simulation, both extracellular and intracellular, are identical to the reference simulation.

Figure 14 presents the changes caused by these oscillations in the environment of the spheroid. Overall, the oscillations generate the same trends as the reference simulation in the evolution of depletion fronts for oxygen and lactate. Whether for 18, 36 or 72 hour oscillations, these depletion fronts progress in the same way, which underlies a similar growth of the spheroid. However, the concentrations that define them do not have the same values. Oxygen being imposed cyclically, it of course alternates between its variation limits, lactate accumulates at slightly higher concentrations (between 8 and 10 mM with oscillations, against 6 mM on the reference simulation).

**Figure 14:**
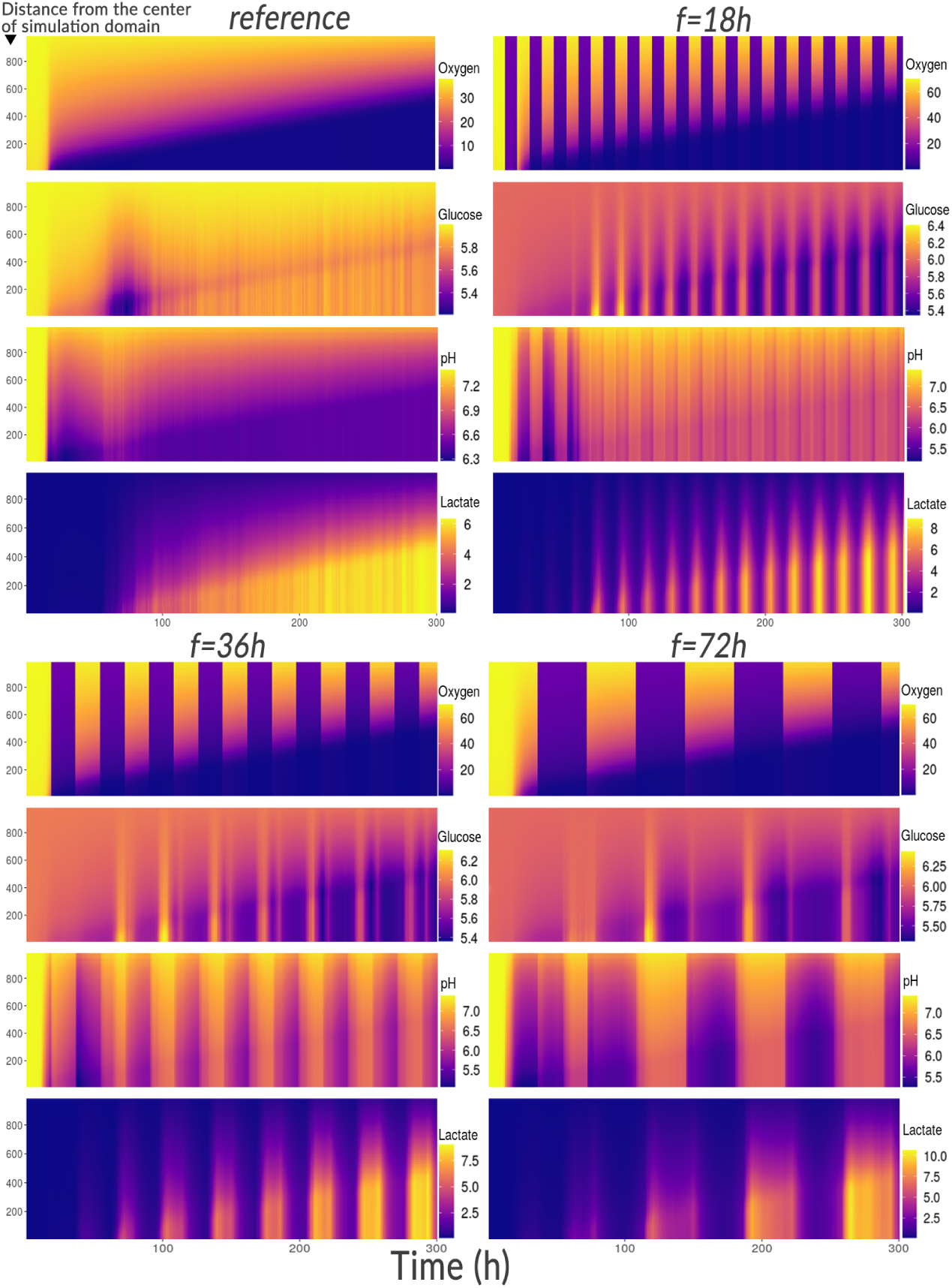
Average radial profiles of substrate concentrations and pH in the environment during hypoxia cycles. The maps presented correspond to three simulations of 300*h* during which cycles of hypoxia with a frequency of 18, 36 and 72h are imposed in the environment. The ordinate shows the distance (in *µm*) from the center of the simulation domain and the color code indicates the extracellular concentrations (mmHg for oxygen, mM for glucose and lactate and without units for pH).

These increases in lactate are synchronous with the drops in oxygen and last for the duration of these hypoxic periods. As for the pH, it does not quite follow the same profile as the reference simulation. Also synchronous with the drops in oxygen, the pH however increases again when the oxygen decreases. Moreover, the lower the frequency of oscillation, the greater the decrease in pH which is further maintained over time. This appears to be inconsistent with cell metabolism. While lactate increases when oxygen is decreased (a sign of increased fermentation), extracellular pH should also decrease (lactate normally exits the cell only through proton excretion).

To understand what is happening, the glucose profile gives indications. When the oxygen in the environment is reduced, the glucose after a short time (about 6*h*) seems to increase suddenly (sometimes beyond the concentration at the boundaries) to decrease again for the rest of the hypoxic phase (intensification of fermentation). What this momentary rise in glucose shows is the synchronous death of a certain quantity of cells which release their contents into the environment. Since the cells are less acidic than the outside, their death increases the extracellular pH and also increases the concentration of lactate and glucose. When the cells again benefit from oxygen, the pH decreases and the lactate also a little later.

The phenotypes associated with these variations are now evaluated. Figure 15 traces over time the proportion of each phenotype in the population of cells still alive. For this, the number of cells associated with each phenotype is divided by the number of living cells. At the start of the simulation, cell metabolism stabilizes, leading to visible variations both in the reference simulation and in the others. When the cells are initiated at the start of the simulation, their internal concentrations do not quite correspond to the ideal concentrations imposed by the environmental constraints. They therefore need a few hours to converge on the nearest metabolic attractors. The addition of oxygen oscillations slightly delays this process but the metabolic behavior stabilizes quickly and is clearly identifiable after about fifty hours.

**Figure 15:**
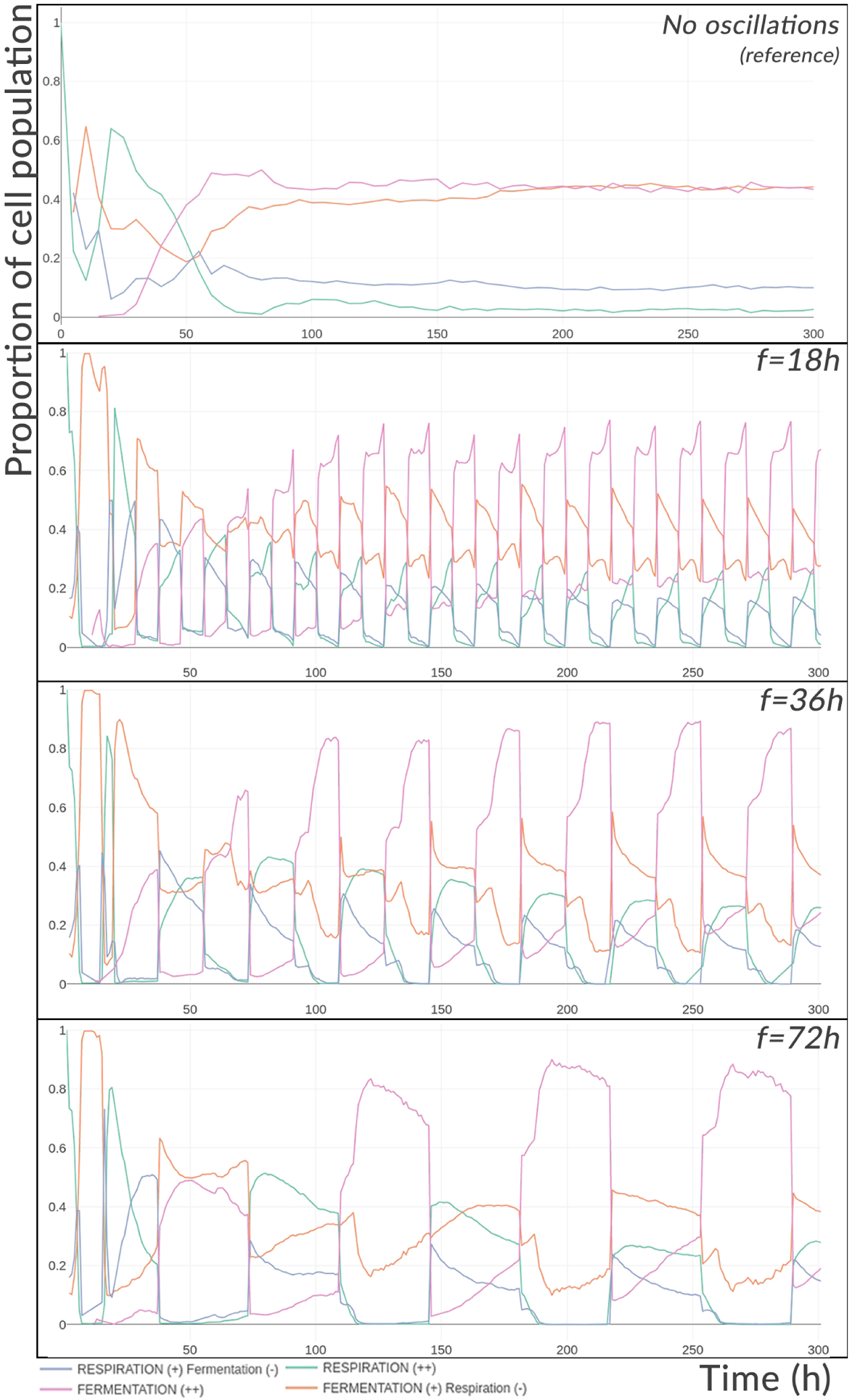
Evolution of the proportion of metabolic phenotypes in the spheroid with or without hypoxia cycles. When a phenotype is described with a (-) it means that it is present but in a lower proportion than that marked with a (+).

Thus, fermentation is the dominant phenotype of the cell population in the hypoxic period with between 75% (for the highest oscillation frequency) and 90% (for the lowest oscillation frequency) utilization. Conversely, the “Fermentation (+) Respiration (-)” hybrid mode is predominant during the oxygenation period (between 40% and 55%). We can note a progressive fall in the use of respiration (majority at the start during the oxygenation period) within the population whatever the simulation considered, due to the increase in the size of the tissue and the consumption of more significant amount of oxygen through the resulting tissue. It should however be kept in mind that this segmentation of the phenotypes serves above all to categorize cell groups and the transitions between states being continuous, the state presented as the majority “Fermentation (+) Respiration (-)” using a little more fermentation than respiration is not much different from the “Respiration (+) Fermentation (-)” state. It is interesting to note that the distribution of cellular energy phenotypes practically does not vary from one cycle to another. This conservation of the oscillatory behavior shows the capacity of adaptation of the metabolism (at least in the short term) to the variation of oxygen.

The translation of these phenotypes by the Warburg and reverse Warburg effect is presented in figure 16. The first frame recalls the distribution of the expression of the Warburg effect in the reference simulation. About 18% of the cells express the Warburg effect (according to the modern definition corresponding to the excretion of lactate in a situation of hypoxia). If we focus on the other cells excreting lactate (excluding hypoxia), the proportion stabilizes at around 80% and represents the majority of the tissue. Thus 98% of the cells excrete lactate. In the simulations of oxygen oscillations, the trends of the reference simulation smoothed over time are still globally preserved. However, locally the oscillations make it possible to greatly vary the expression of the Warburg effect. The lower the oscillation frequency, the higher the amplitude of variation in expression of the Warburg effect. When the cycles are long, the cells have more time to converge on a metabolic attractor and therefore generate more marked variations during changes.

**Figure 16:**
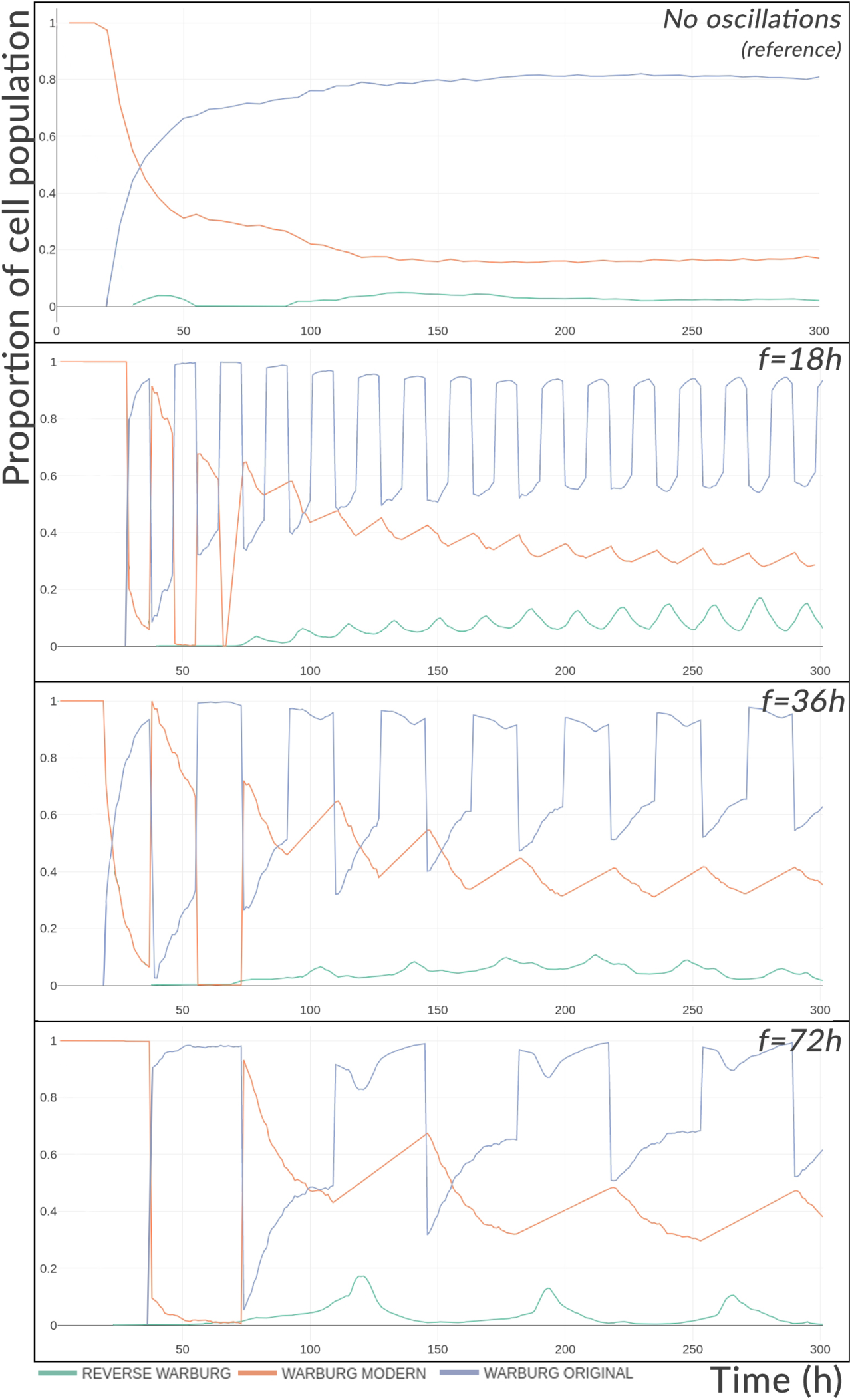
Evolution of the proportion of Warburg and reverse Warburg effect in the spheroid with or without hypoxia cycles.

The 72-hour cycle in particular, creating longer fermentation times, generates more lactate but also has more time to let the reverse Warburg effect occur during the reoxygenation phases and return to its original level. On the contrary, the shorter cycle of 18 h, creates a little less lactate but sees the basal activity of the reverse Warburg slightly increase, the higher frequency of oscillation leaving less time for the cell to equilibrate its intracellular concentration and extracellular lactate. Thus the reverse Warburg effect can represent at certain times, 19% of the metabolic expression of the tissue, *i.e.* as much as the basal expression of the Warburg effect (modern definition) of the reference simulation.

#### 3.2.2 Acid shock

The spheroids are now subjected to sudden drops in extracellular pH in order to test the metabolic consequences of sudden exposure to the acidic environment encountered within tumors. For this, the entire environment (not just the domain boundaries) is maintained at physiological pH (= 7.4) for 50*h* then, from the fiftieth hour, it is maintained at pH values lower than or equal to the physiological pH.

Figure 17, presents the results of these simulations. The first frame recalls the reference simulation in which the pH is only imposed at the domain boundaries and evolves freely in the rest of the simulation domain. The second frame presents the four pH tested: 7.4 (the pH is not changed and is maintained throughout the environment at this physiological value), 6.7, 6.0 and 5.3. The metabolic landscape is plotted before the change in acidity at 50*h* and after at 130*h*. Here there are no oscillations and the metabolic landscape converges to those presented.

**Figure 17:**
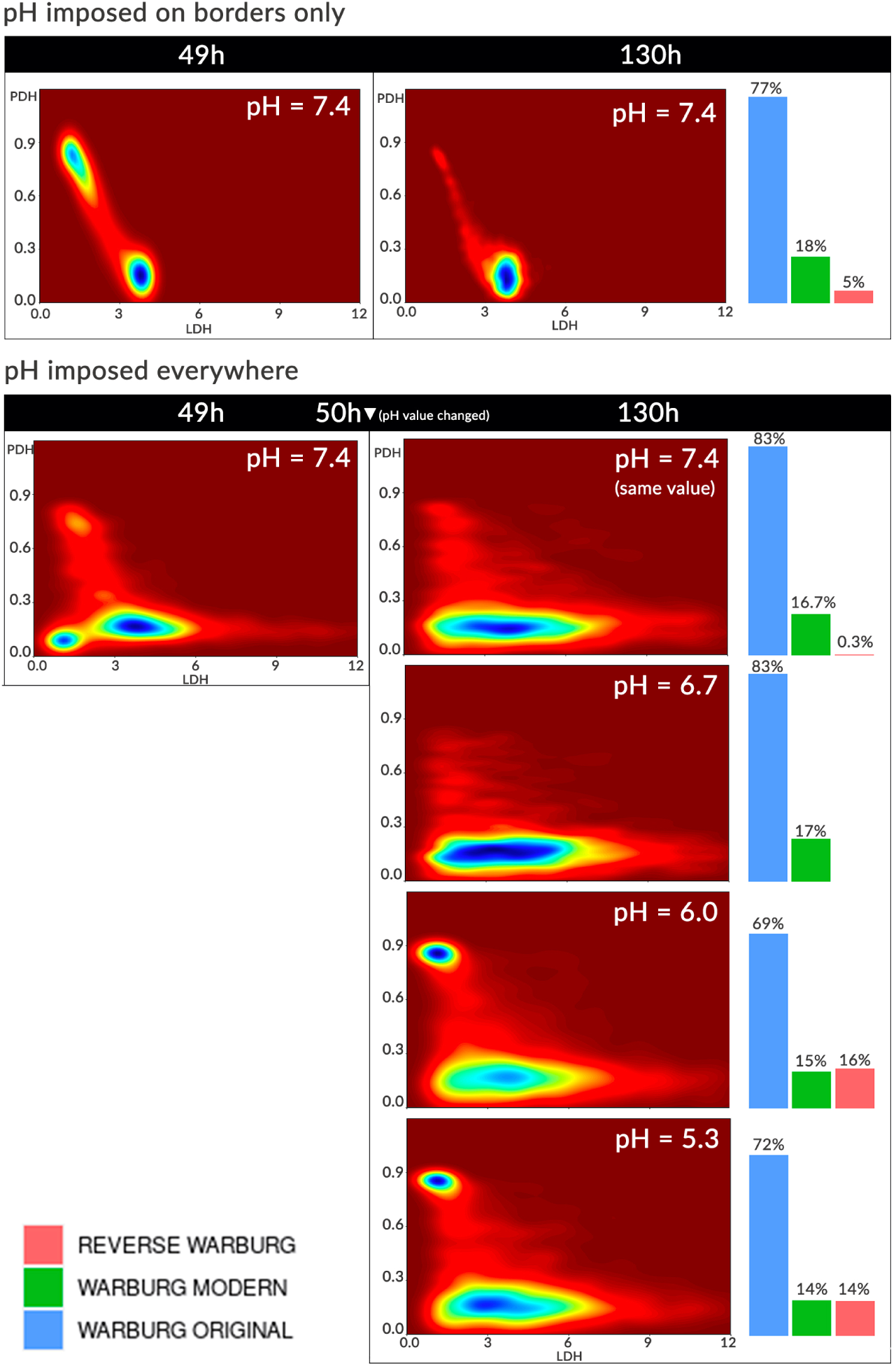
Metabolic landscapes and expression of the Warburg effect before and after pH change in the whole environment. The “pH imposed only on the domain boundaries” situation corresponds to the reference simulation.

The first observation that can be made before the pH changes, is the difference between the environment with pH imposed at the domain boundaries and that with pH imposed everywhere. When the pH is only imposed at the boundaries, the pH becomes increasingly acidic at the core of the spheroid as glucose is consumed and lactate excreted along with protons. This reduces the fermentation capacity of the cells which are mainly located in the *intermediate* zone of the landscape (defined around LDH ≈ 4 and PDH ≈ 0.1). A part of the cells (those with more access to oxygen) continue to use respiration a little more in the oxidative zone of the landscape (0.9, 1.0). On the contrary, when the pH is imposed everywhere in the environment, the cells mainly in the *intermediate* state, also cover a larger space of the metabolic landscape and suggest a distribution between its four attractors (*neutral, intermediate, oxidative* and *glycolytic*). The glycolytic state is the least expressed at the landscape level but is not absent.

At 130*h*, the reference simulation indicates that the majority of the cells have centralized in the *intermediate* state, the few cells benefiting from a little more oxygen gradually ending up transiting towards this state as the tissue grows and the oxygen available per cell decreases. This behavior is found again at 130*h* and at fixed pH with, however, a widening of the *intermediate* zone. The cells can be more glycolytic, the acidity being either less strong (when the pH remains between 6.7 and 7.4) or established for less time (since only *t* = 50*h*) than the reference simulation. Going down in acidity however, between pH 6.7 and 6.0, (visible in landscapes from pH 6.0), a focus of cells in the oxidative zone clearly re-emerges, the pH here being low enough for certain cells to inhibit glycolysis and the amount of accumulated lactate sufficient to introduce the reverse Warburg effect. It is from these pH that the reverse Warburg effect reaches the same proportion (approximately 15%) as cells excreting lactate when oxygen is insufficient (Warburg effect by its modern definition).

Here the pH has a double effect. By being identical everywhere, it initially allows cells to explore a wider and more permissive metabolic zone. The reachability of these areas is then directly linked to access to the main substrates, which are oxygen and glucose. Secondly, at a more acidic pH, it inhibits glycolysis. Whether this is through some form of negative feedback (in the case where fermentation is responsible for the pH drop), or whether it comes from an imposition of pH in the environment, the result is the same, fermentation is impaired.

This is also what figure 18 shows. When the pH is maintained throughout the environment and particularly at pH 7.4, fermentation is maintained more easily than in the reference simulation, acidity not inhibiting glycolysis. Before 50*h*, the distribution of the phenotypes is indeed identical and deviates after the change in pH, mainly for the most acidic pHs (6.0 and 5.3). At pH 6.7, a very slight drop in the proportion of fermentative cells is observed compared to the pH maintained at 7.4, and rises again to the same level as this. This trajectory for fermentative metabolism is associated with the opposite trajectory for respiratory metabolism, which increases very slightly and then decreases. At pH 6.0 and 5.3, respiration and the “Respiration (+) Fermentation (-)” hybrid phenotype are more greatly increased, with the respiration rate in the tissue reaching almost the same rate as the “Fermentation (+) Respiration (-)” at 60*h*. At the same time, the proportion of essentially respiratory cells drops by around 35%.

**Figure 18:**
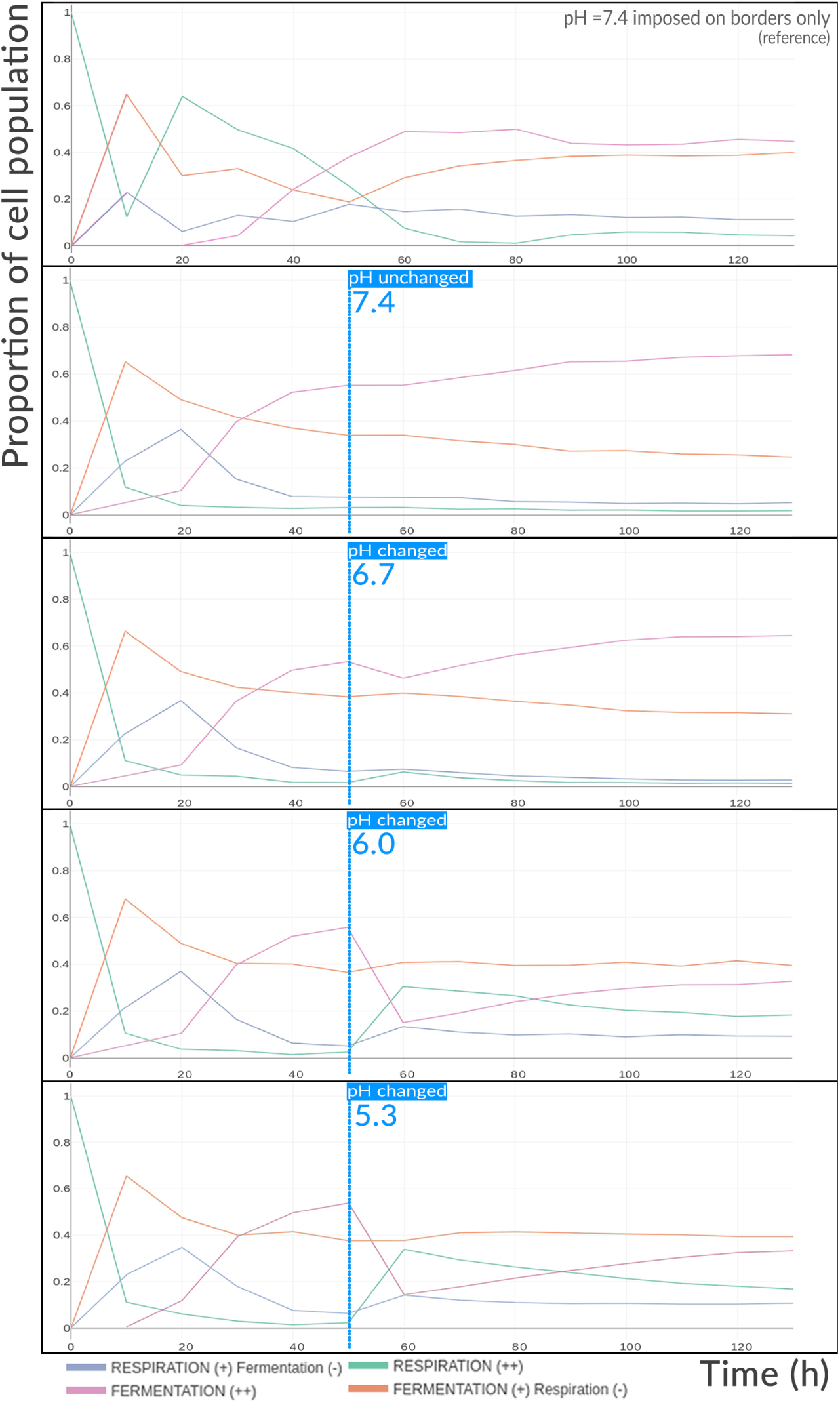
Evolution of the proportion of metabolic phenotypes in the spheroid population over time before and after the pH change in the whole environment.

Then over the following hours, the distribution of the different phenotypes tends to reverse again. Fermentation increases and respiration decreases, without however reaching the rates observed at physiological pH. The reason for this rebound is the same for the figure 17: the progressive drop in oxygen accessible by the cells of the deeper layers of the growing spheroid. Glucose is here in excess compared to the quantity of oxygen available and the cells have the advantage of storing glucose. In the absence of oxygen, and despite acidic pH, fermentation increases but with reduced activity.

#### 3.2.3 Glucose depletion

In the same way as was done for pH, the following simulation aims to look at the consequences of glucose depletion on the tumor. In the model, apart from oxygen, glucose (and lactate) are the only external sources to replenish pyruvate or acetyl-CoA levels in the absence of fatty acid or glutamine. The idea in this simulation is to look at the evolution of the metabolism when deprived of its main source of carbon.

The spheroid of this new simulation grows under the same conditions as the reference simulation with, however, the glucose concentration fixed at 6.0 mM in the whole environment (not only at the domain boundaries). Since the glucose gradient introduced into the reference simulation before 60*h* is negligible, it is possible to maintain the glucose at a constant concentration before. After 60*h*, the glucose concentration is reduced to 0.01 mM throughout the environment and maintained as such. This fall is visible on the figure 19. The simulation lasts 300*h* in total. It is longer than the pH simulation presented above, because the effects of glucose depletion take longer to emerge.

**Figure 19:**
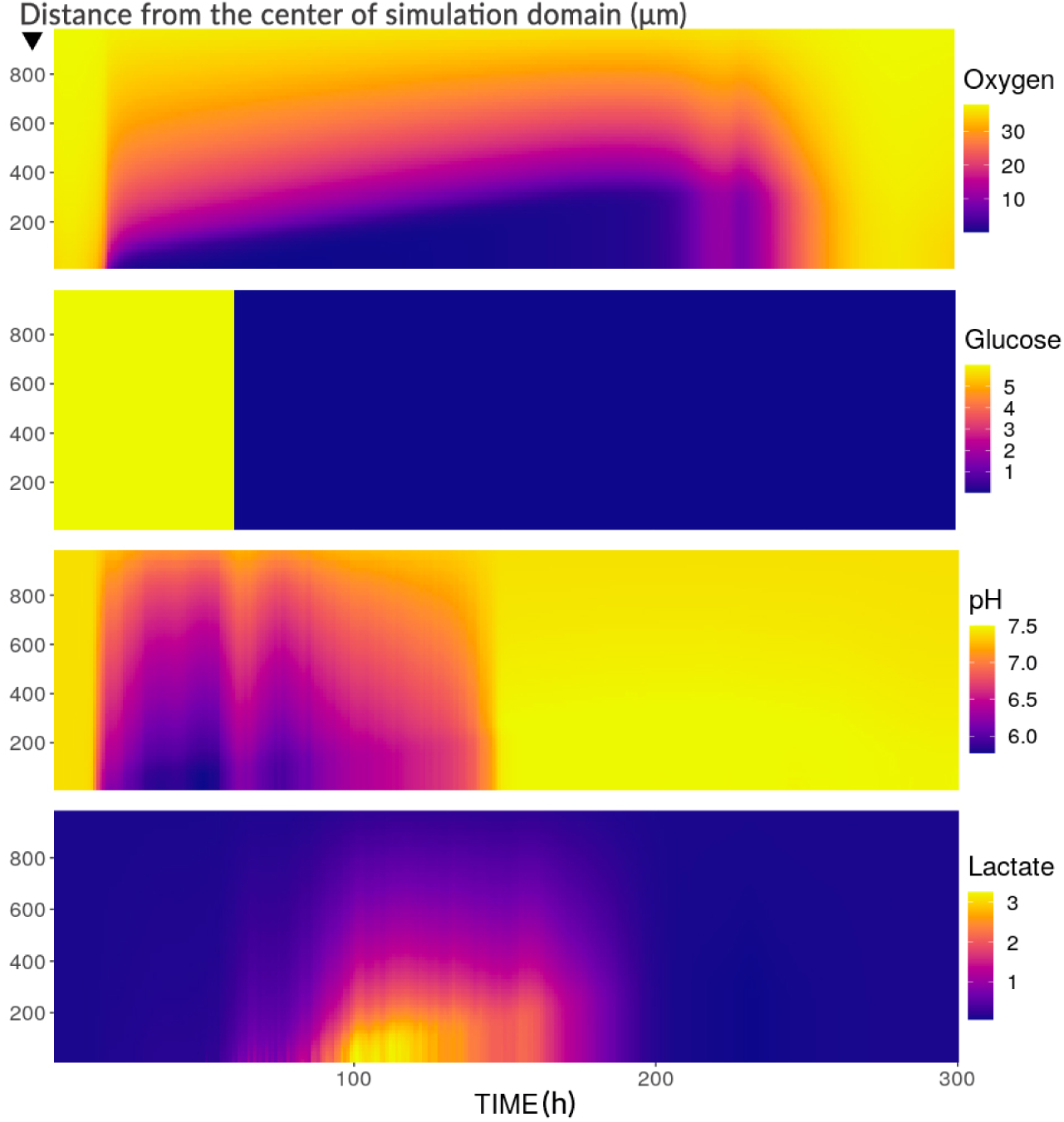
Mean radial profiles of environmental substrate concentrations before and after glucose depletion at. 60*h*. The ordinate shows the distance from the center of the simulation domain and the color code indicates the extracellular concentrations (mmHg for oxygen, mM for glucose and lactate and without unit for pH).

Unlike previous simulations, here for oxygen, pH and lactate, the progression of their radial footprint is not linear. The acidic pH becomes physiological again around 150*h*. The lactate reaches its highest concentration at 100*h* (approximately 3 mM) and drops to almost 0 mM shortly before at 200*h*. Oxygen depletion progresses as before until 230*h*, and after a small re-increase in its concentration immediately followed around 240*h* by a short episode of depletion, oxygen returns durably to its original concentration of 38 mmHg.

Figure 20 sheds some light on what is going on. Cell states and cell counts hardly deviate from those of the baseline simulation until 150*h*. It is from this moment that there is a drop in the number of proliferating and quiescent cells. This decrease continues for the proliferating cells up to 200*h*, in favor of the number of quiescent cells which increases. The number of dead cells temporarily stabilizes and increases again around 230*h*, while the number of quiescent cells decreases rapidly. The burst of oxygen observed between 230 and 240*h* on figure 19 corresponds to this decrease in quiescent cells combined with the very slight and momentary increase in the number of proliferating cells which then decreases until the end of the simulation leaving only a hundred cells still alive.

**Figure 20:**
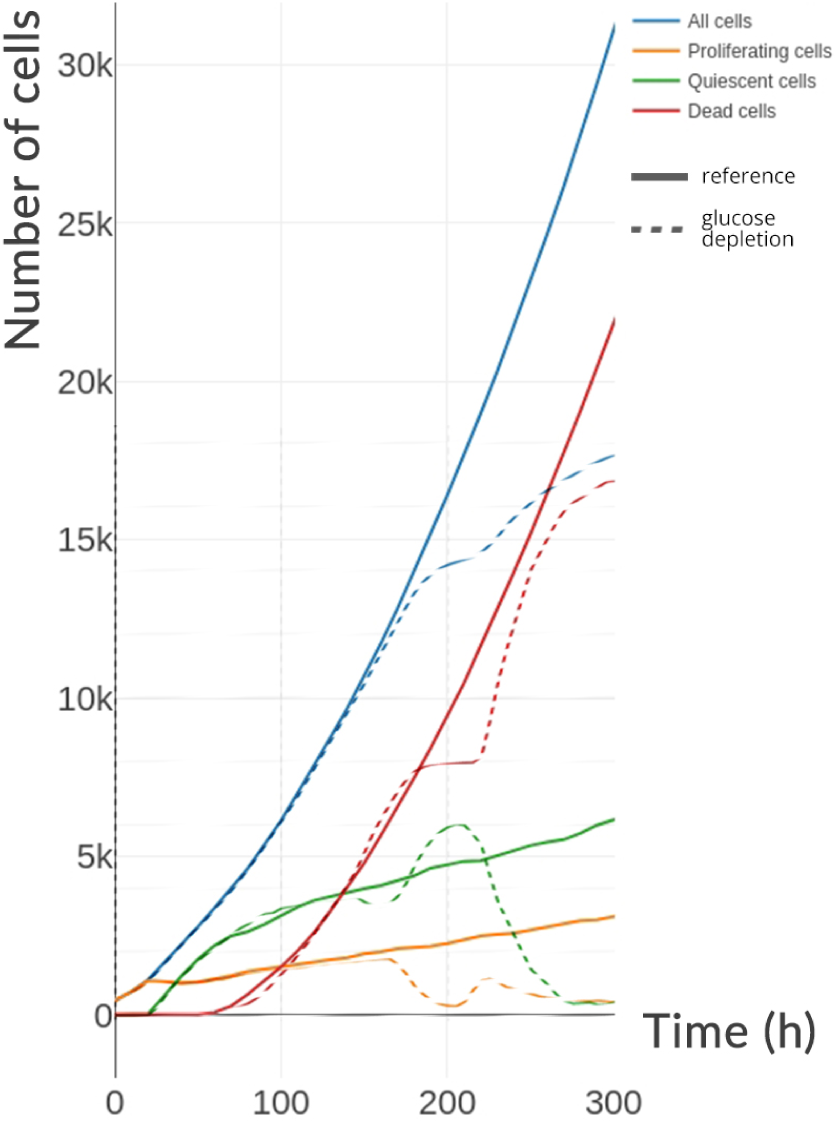
Evolution of cell states within the population of spheroids over time before and after the drop in glucose in the whole environment. The simulation of the drop in glucose (dashed lines) is compared with the reference simulation (solid lines). Glucose depletion takes place from 60*h*.

Thus from the point of view of tumor growth, the tissue was not disturbed immediately after the drop in glucose, the disturbance only occurred 80 hours later. The reason stems from the delay between glucose depletion in the environment and intracellular depletion. Where intracellular oxygen pressure equilibrates with extracellular pressure by simple diffusion, intracellular glucose concentration equilibrates only by facilitated diffusion [29] which, although faster than simple diffusion, depends on the number of GLUT membrane transporters. There is therefore a time during which the glucose leaves the cell (without increasing the extracellular concentration, the latter being kept fixed at each time step) and is at the same time consumed by the same cell through glycolysis. This is a kind of time to exhaust its reserves.

Then, as intracellular glucose falls, pyruvate is produced less and less by glycolysis but its concentration must be maintained to maintain sufficient energy production. The lactate produced until then by fermentation is also consumed. Its intracellular depletion also results both from its rate of consumption (transformation into pyruvate) and from the depletion of lactate in the environment. Thus, a little before 200*h*, almost all of the extracellular lactate has been reintroduced into the cells, causing at the same time a rise in the pH and from 210*h* onwards the intracellular lactate in the majority of the cells is consumed (not shown). Respiration can then no longer take place and the oxygen which is no longer used by the cells gradually rises. Cells that no longer produce enough ATP mostly die. There are still a few cells still alive which suddenly benefit from ideal oxygenation, the little glucose still present and the death of neighboring cells which release the remains of glucose and lactate from their cytoplasm into the environment.

Figure 21 confirms delays in metabolic responses to glucose depletion. Before 60*h*, the phenotypes and the expression of the Warburg effect are almost identical to the reference simulation (Fig.16) with however a slightly greater increase in the reverse Warburg effect due to the amount of lactate accumulating more rapidly within the cells (due to the absence of a gradient). After 60*h*, the proportions do not really change apart from the fermentation (“Fermentation (++)”) which begins to decrease at the same time as the intracellular glucose and the reverse Warburg effect which continues to increase.

**Figure 21:**
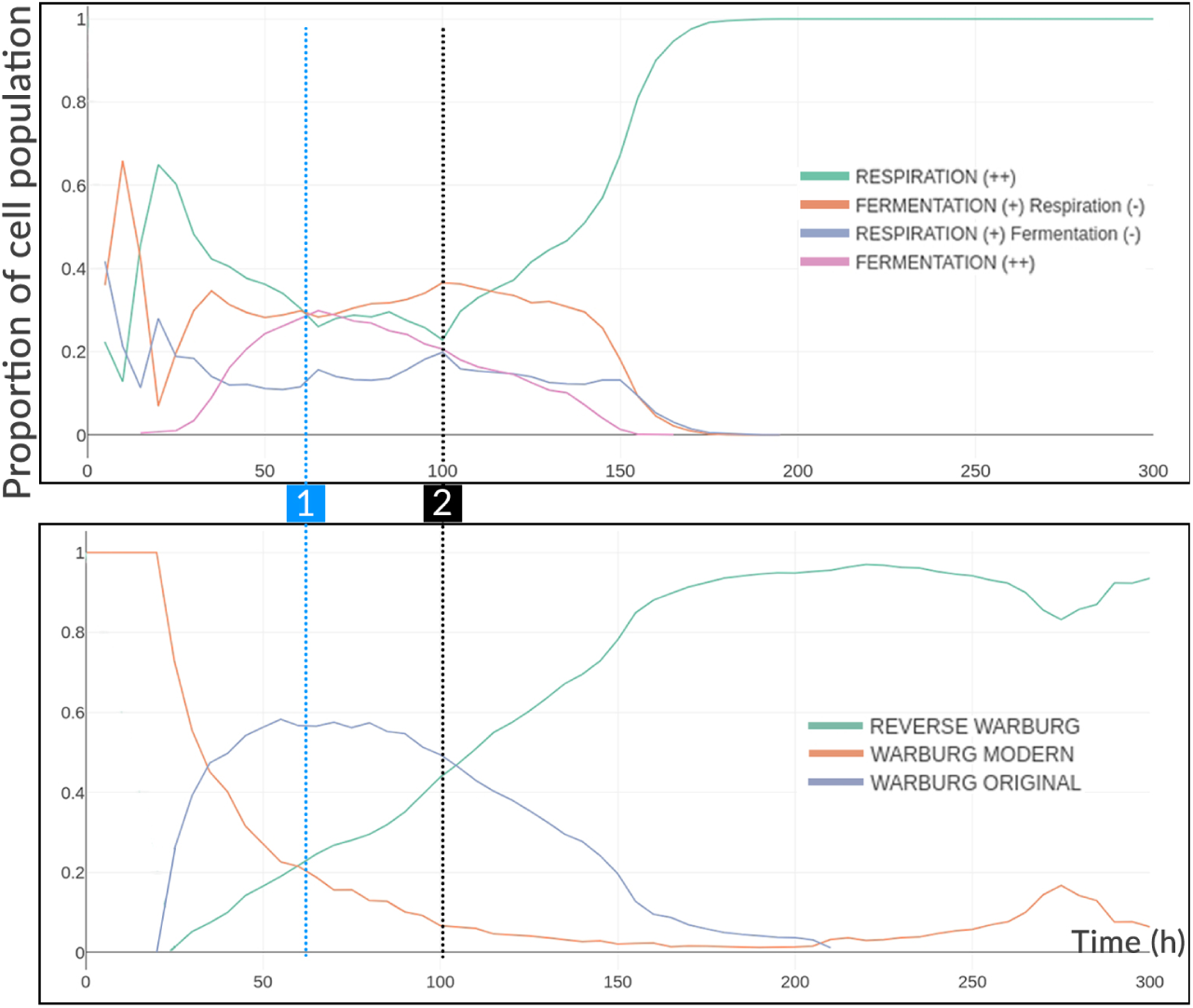
Evolution of the proportion of metabolic phenotypes and expression of the Warburg effect in the cell population over time before and after the drop in glucose concentration in the whole environment. 1) At 60*h*, the extracellular glucose is fixed throughout the environment at 0.01 mM. 2) At 100*h*, the intracellular glucose of the cells has reached the extracellular concentration.

From 100*h*, when the concentration of intracellular glucose has reached the extracellular concentration, the rate of respiration slows down and begins to increase to become at 180*h* the almost unique mode of functioning of the cells. From the perspective of the Warburg effect, it persists longer than fermentation does, with intracellular lactate levels continuing to equilibrate externally, with lactate sometimes being excreted. This also shows the fact that the Warburg effect is not always induced by the concomitant fermentation activity.

### 3.3 Challenging the glycolytic metabolism

The intrinsic heterogeneity of cell metabolism makes it difficult to derive a general law for tissue evolution in response to environmental conditions. In particular, tissue dynamics quickly take precedence over the individual metabolic dynamics of each cell. Thus, on many simulations, the behaviors that emerge are not always systematically anticipated. However, from a large number of simulations that we realized the following trends emerge:

Thus, lactate is generally excreted from the cell when oxygen is in low quantity in the environment or when lactate is already present in the environment. Indeed, when lactate is present, the expression of the HIF-1α gene is increased, which induces (although there is oxygen) an increase in glycolysis. If there is no lactate but glucose and oxygen in sufficient concentrations, then the production is generally directed towards respiration, the production of lactate is relatively low (but not non-existent).

On the other hand, if lactate is present in large quantities in the environment, the cells will tend to import it, following the concentration gradient. However, this does not prevent the cell from continuing to produce it (by fermentation) by accumulating it inside. Lactate is also imported regardless of its concentration, in the presence of oxygen and combined with the absence of glucose, to replenish the level of pyruvate used by the mitochondria.

What makes it difficult to identify these trends comes from the sources of variability in the conditions external to the cells. Fixing the levels of oxygen, glucose, lactate or the pH, does not prevent the cells of a layer of the spheroid from generating locally, contrary situations. For example, cells in hypoxia produce lactate in large quantities and lower the pH of cells at the periphery which find themselves forced to use lactate instead of glucose. These interactions between cells are an important source of heterogeneity often masked by the averaged behaviors evaluated in a good number of experiments [30].

To illustrate this, a new simulation is presented. In this simulation, the oxygen is at atmospheric pressure with 160 mmHg *i.e.* in very great excess, whereas the glucose is only at 1 mM, which is a very small quantity. These values are kept constant at the domain boundaries. The lactate at the start of the simulation is at 0 mM. The initial pH is acidic at 6.7. The boundary conditions for lactate and protons are Neumann’s conditions (zero flux) which means that lactate can accumulate without being eliminated by the medium during the course of the simulation and the same for the protons. Cells can only use very little glucose, because it is present in very small quantities and the initially acidic pH slows down glycolysis.

The objective of this simulation is to assess whether tumor cells really remain in a glycolytic metabolism when the conditions for this are unfavorable. In other words, is the Warburg effect (with the modern definition, excretion of lactate in the presence of oxygen) systematic? Previous simulations have already shown that this was not the case, however this simulation is also interesting from the point of view of cellular interactions.

Figure 22, shows quite logically, that the tissue uses pyruvate through respiration (dark purple cells). Where oxygen is rarer in the center, some cell islands have no choice but to use the little glucose available. The cells are indicated for a large part of them as having a reverse Warburg type phenotype, while a vertical and centered bundle of cells produces the Warburg effect. Again, fairly large areas excrete lactate while having a predominantly oxidative phenotype. Glucose, although in very small quantities, is continuously available from the edges and lactate can accumulate (Fig.23).

**Figure 22:**
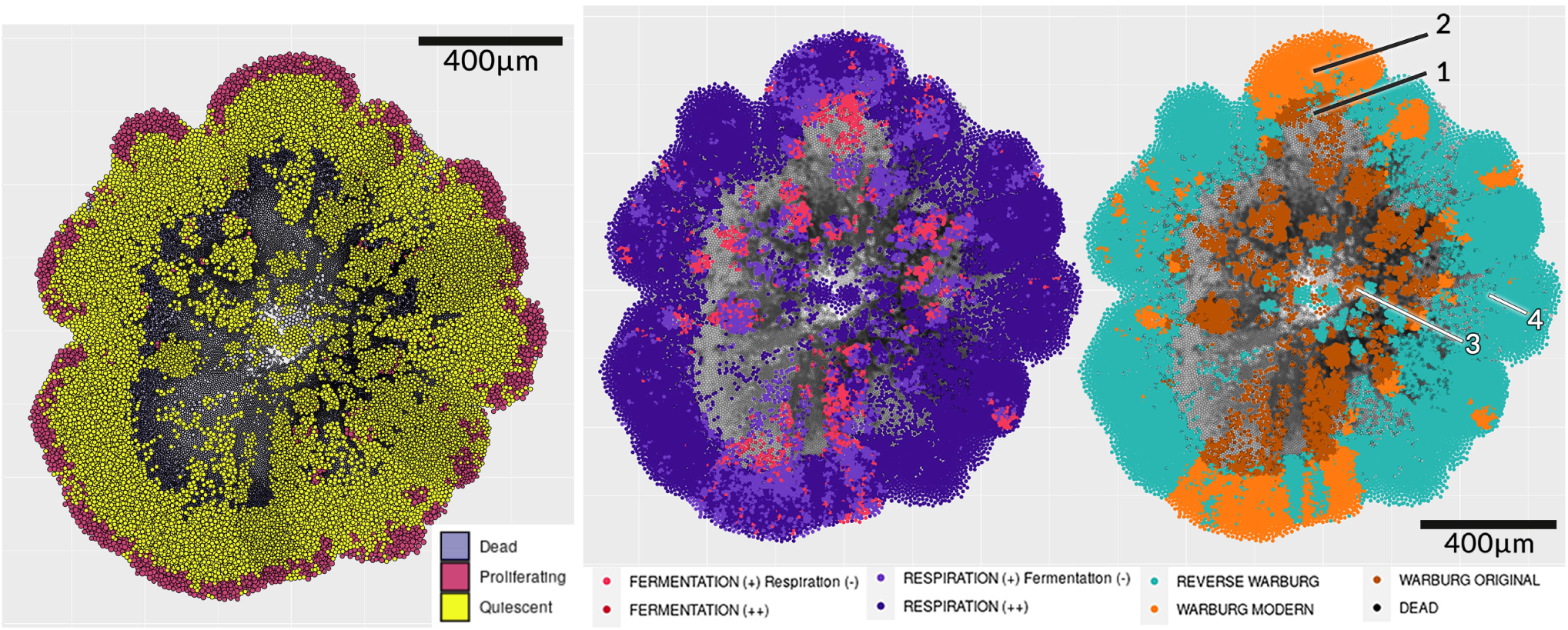
Cell phenotypes of the spheroid at 420h. On the left the cell states. In the center, the staining corresponds to cell phenotypes of fermentation and/or respiration. On the right the cells are stained in relation to their phenotype corresponding to the definitions associated with the Warburg effect. When a process is described with a (-) it means that it is present but in a lower proportion than that marked with a (+). The black cells are in necrosis.

**Figure 23:**
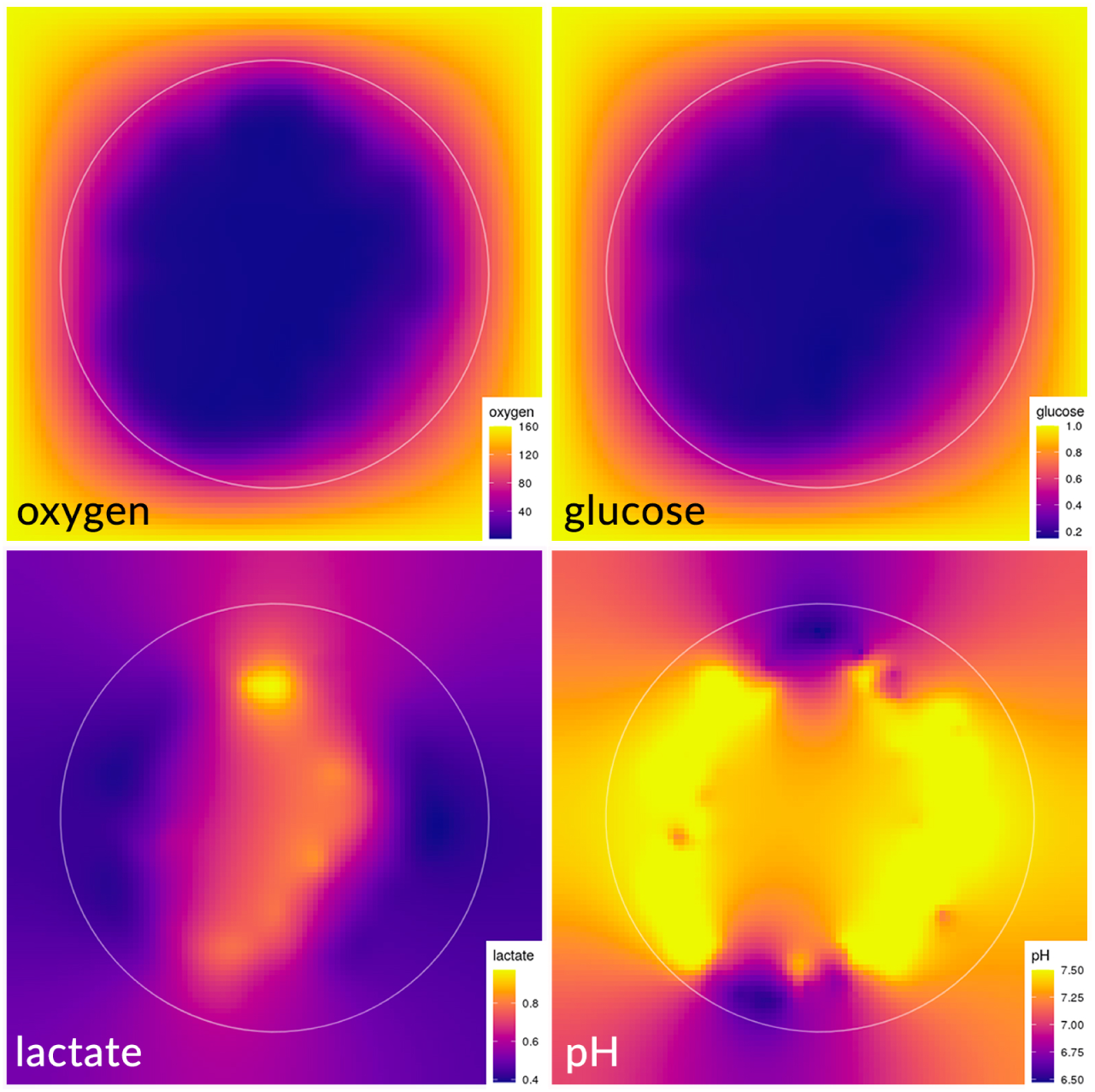
Extracellular concentrations of the substrates at 420h. Oxygen (top left) concentrations range from 0 to 160 mmHg, glucose (top right) from 0.2 to 1.0 mM, lactate (bottom left) from 0.4 to 0.97 mM and pH (bottom right) from 6.49 to 7.49.

After a certain time, the cells that die (close to zone 3 on figure 22), release lactate, glucose and protons which are quickly imported by the surrounding cells. In an emergent way, cell death has generated an internal asymmetry in the tissue as well as at the level of the diffusion of the substrates. The little glucose consumed by cells in hypoxia (particularly in zone 1) is used to produce lactate and acidity which accumulate in these cells (zone 1 and 2) and in the medium (at the balance between the two).

In doing so they increase their level of pyruvate and more easily maintain their level of ATP. Surrounded by lactic acid, the cells of zone 2 reduce their consumption of glucose and emit the lactate that they have in their cytoplasm and coming from zone 1. Thus the lactate is found outside the spheroid and diffuses into the rest of the environment. This lactate is recovered by the cells on the sides of the spheroid (as in zone 4) which, by transporting it inside their cytoplasm, reduce the pH in the surroundings. Most of the cells on the periphery of the spheroid therefore have a predominantly oxidative metabolism, but in the face of glucose depletion and localized acidity conditions, it is the greed for lactate, a source of potential energy, that creates the tissue dynamic.

Figure 24 confirms this mechanism. Cells in hypoxia (zone 1) appear clearly blue on the spheroid with LDH flux staining. The pyruvate is directed towards the production of lactate whereas for the others in red it is the lactate which is transformed into pyruvate. For lactate and proton fluxes (MCT), at this stage of the simulation, lactate is essentially imported into the cells on the sides of the spheroid.

**Figure 24:**
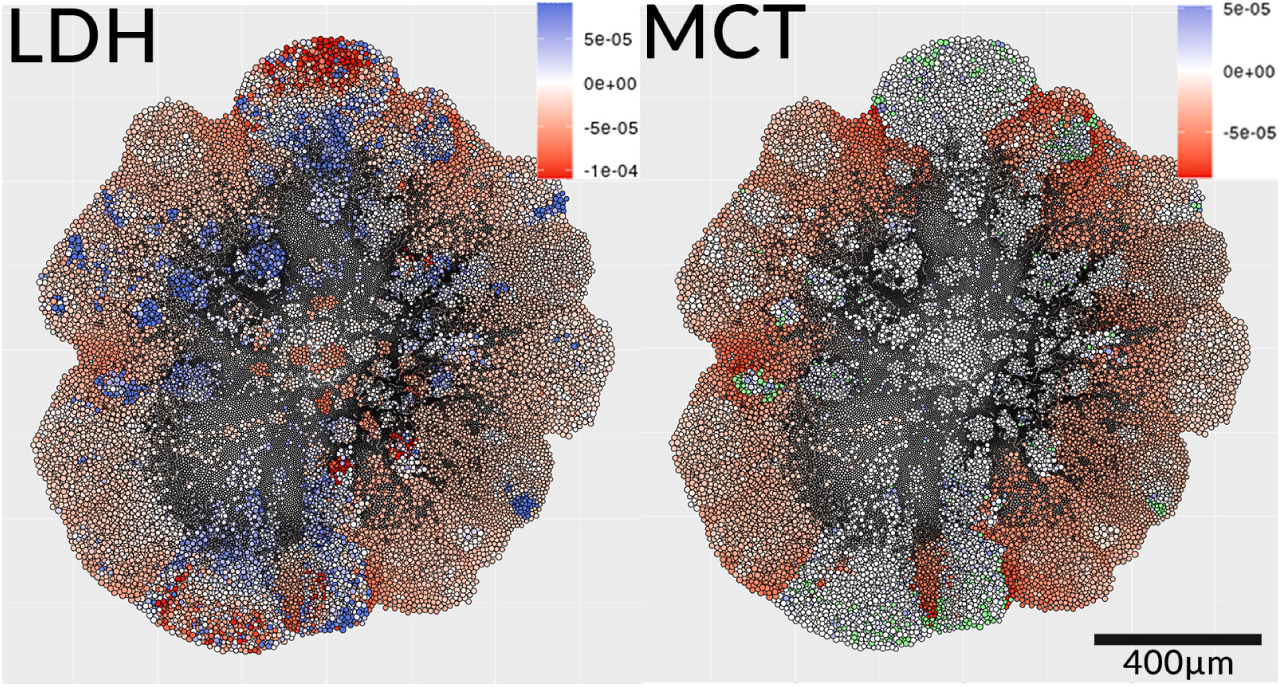
Flow of the reaction catalyzed by LDH (r12) and transport of lactate via MCT channels (r21) in the spheroid at 420h. The fluxes are expressed in mM/min. A negative value indicates that the reaction or transport is in the opposite direction (lactate to pyruvate and extracellular lactate to intracellular lactate). The few cells in green on the right spheroid represent those that produce much more lactate than the others (about 20x more). They appear in green so as not to interfere with the coloring of other cells.

Cells appear mostly white in zone 2 and excrete little lactate (relative to those that import it). Rare cells (in green) having accumulated a lot of pyruvate excrete a lot of lactate (about 20x more than the others at about 10*^−^*^3^ mM/min).

These simulations show that it is uncertain to speak of the Warburg effect when thinking of targeting a single type of metabolism in a particular way. If the Warburg effect appears on a macroscopic scale (at the tissue level) this is not particularly the case at the level of individual cells. As we already explored this question in [1, 2], the definitions attached to it do not make it possible to specify the contours of a precise metabolic behavior at the cellular level and even less of the threshold values (for example *is the Warburg effect the production of X amount of lactate in the presence of Y amount of oxygen*). Even without defining these thresholds, taking the tissue as a whole and simply considering lactate production (Otto Warburg’s original observation) the Warburg effect is not ubiquitous.

At the level of the metabolic landscape, three zones out of the four are identified (Fig.25). Initially (*∼* 10*h*), the few cells benefit from the little glucose present by focusing their metabolism towards the *glycolytic* zone (*i.e.* at the bottom right, poorly expressed PDH and LDH strongly expressed). Quickly (*∼* 70*h*), part of the cells are pushed to adopt a less glycolytic metabolic mode (return to the zone described as *neutral* at the bottom left) mainly because of the acidity that this mode has generated . These cells move along the LDH axis without actually varying in PDH expression. At 100*h*, the lack of glucose settling permanently in certain areas of the spheroid, pushes the cells in the *glycolytic* and *normal* states to converge towards the *oxidative* state (top left). Cells in the *glycolytic* zone undergo glucose deprivation and increased acidity. Instead of passing through the *neutral* zone first, they are pushed directly into the *oxidative* zone. This phenomenon is visible in particular from 130*h* (Fig.26) and until the end of the simulation.

**Figure 25:**
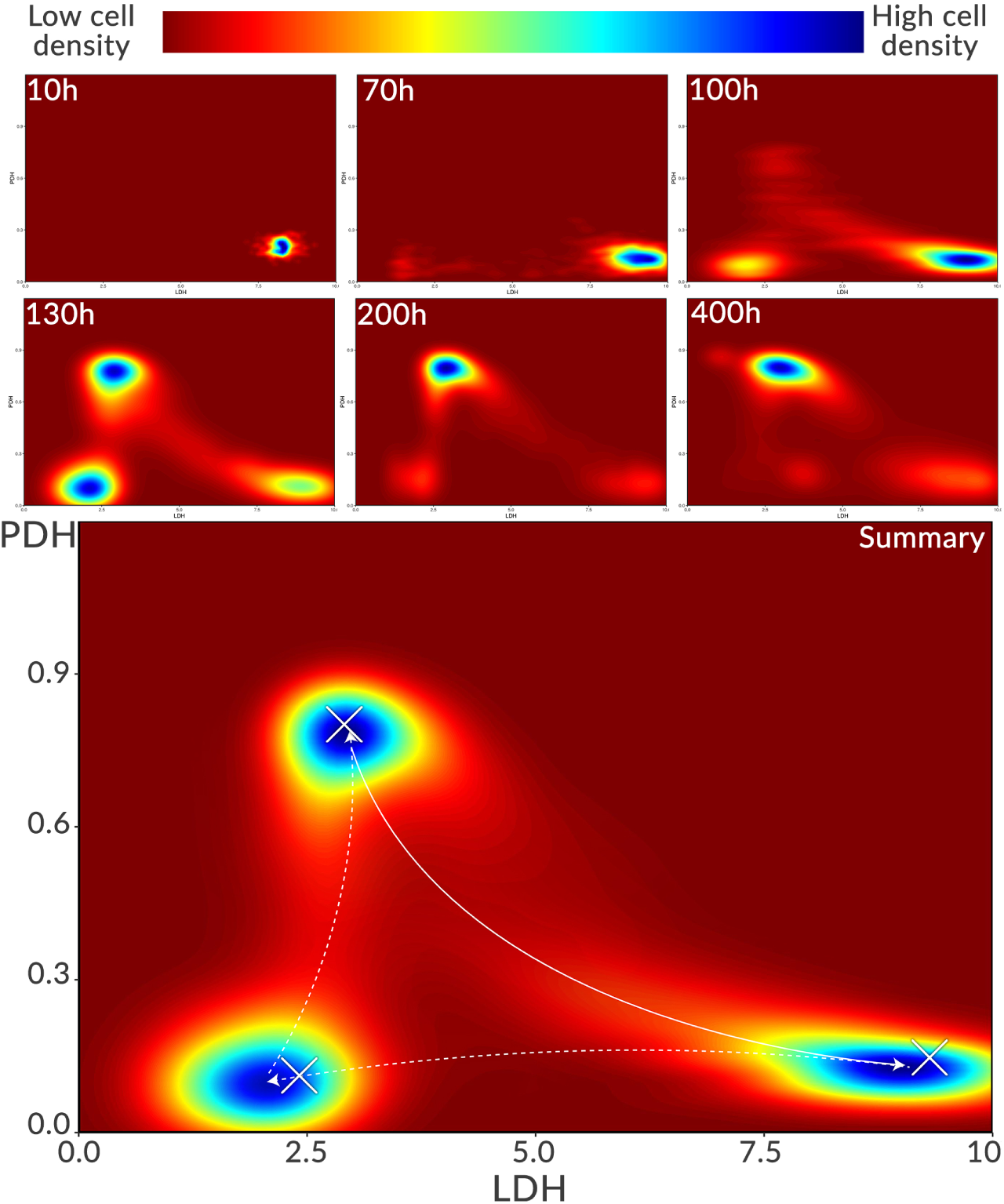
Evolution of the metabolic landscape over time. The maps show the areas of metabolic densities (number of cells), at the level of expression of the LDH and PDH genes. The red areas are areas where very few cells are found, unlike the blue areas which contain a large quantity. The map shown below is the aggregation of the metabolic landscape over time, accumulating areas of high densities. The crosses indicate the regrouping point of the cells.

**Figure 26:**
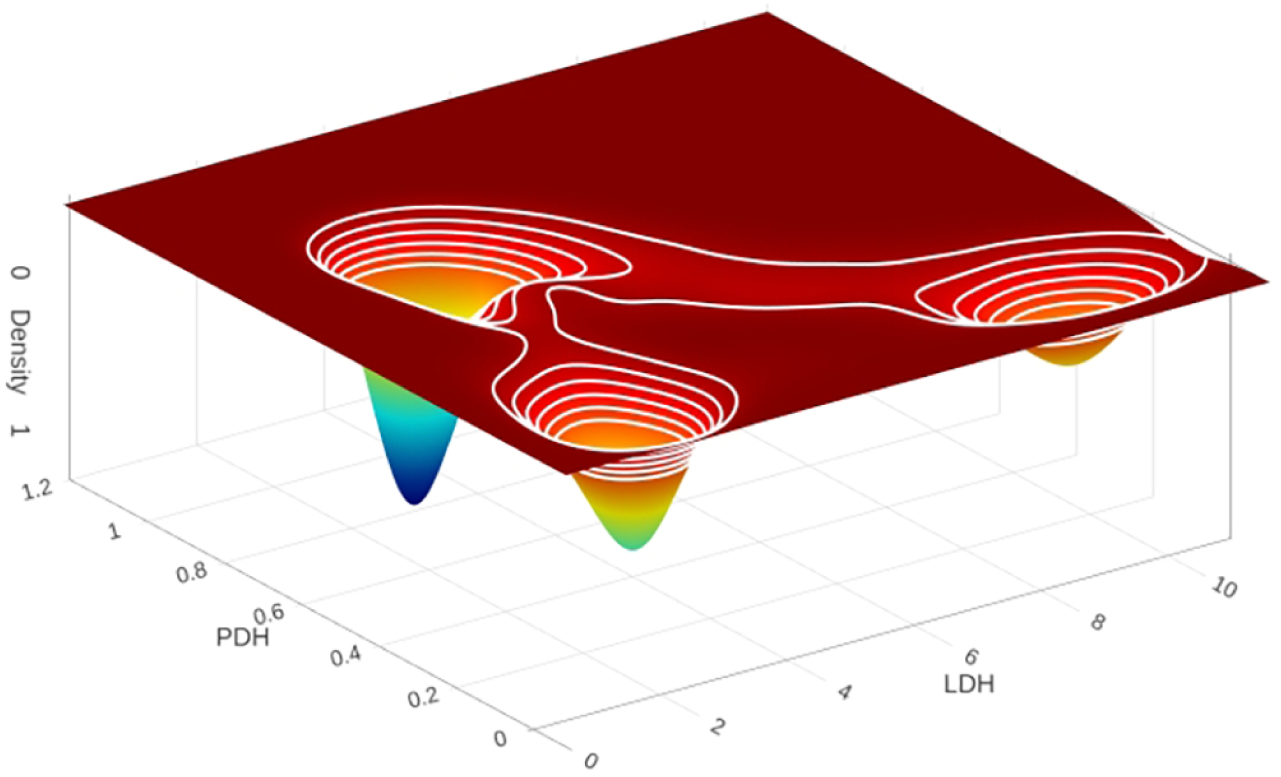
Cell density in the LDH/PDH bifurcation at 130h.

At 400*h*, the cell population is mainly distributed between an *oxidative* state and a *glycolytic* state, but this is an overall view and although the majority of cells are grouped into clusters in these attractors, transitions between the three states continue to arise (visible by the traces between clusters). The last image of the figure 25 represents the aggregation of these three attractors during the simulation and the trajectories described.

## 4 Conclusion

The simulations carried out made it possible to identify in an emergent way, different metabolic behaviors which are spatiotemporally defined. The reference simulation showed the standard radial distribution of cell states (proliferative, quiescent, and necrotic) within a spheroid. It showed the impact of the underlying metabolism on the environment with mainly a strong oxygen depletion in the internal layers of the spheroid and an increase in lactic acidosis. These profiles correspond to classical observations of spheroids *in vitro* [31, 32]. It also highlighted a partial decoupling between the Warburg effect and the type of metabolic phenotype. Essentially respiratory cells can express the Warburg effect by secreting lactate. This results in a marked heterogeneity of the distribution of the cells of the spheroid within the metabolic landscape defined by the expression of the LDH and PDH genes. Ultimately, however, an intermediate metabolic state predominates in the tissue.

Simulations of intermittent oxygen concentration have shown the short-term stability of metabolism to these variations as well as the resurgence of the reverse Warburg effect on tissue reoxygenation phases. The acid shock simulations also showed the stability of the metabolism in the face of sudden pH variations. More acidic pH allow the reverse Warburg effect to be more strongly expressed (phenomenon experimentally observed [6]) which is also reflected in the metabolic landscape by an increase in the expression of PDH. Depletion of extracellular glucose has a slower effect, depending on intracellular reserves. The metabolic impact is limited as long as lactate (and by extension other sources to continue to fuel respiration) is available. Once these sources are exhausted, this condition is most critical for the survival of rapidly dying cells.

The simulations highlight the complexity of the multiple behaviors that can take place within the same tissue, with cyclic hypoxia phenomena when the external conditions are at the limit barely sufficient to allow a few cells to proliferate. There can also be phenomena of indirect cellular collaborations, of “metabolic symbiosis” [33], the metabolic products of a cell exposed locally to certain environmental conditions benefiting another cell exposed to different conditions.

As a consequence, it is possible to conclude that the metabolism is relatively robust in the face of environmental disturbances, at least over short periods of time (<1 month). In reality, what introduces instability in the metabolic organization of a tissue does not come intrinsically from the metabolism, but from the existence of abnormal conditions in which the cells are pushed (the depletion of glucose in the environment resulting in death is an example). A spheroid, too, is an abnormal tissue configuration due to its high density or the absence of vascularization. The metabolism itself has all the biochemical and thermodynamic properties [34] to continue to be operational from the point of view of maintaining cell viability.

## 5 Supporting information

**S1 Appendix. Equations of the model of cell metabolism**. The model for cell energy metabolism by Li and Wang (2020) has been modified as described in the *Models* section. The model comprises 56 ODEs, 119 simple parameters and 3 matrices *S*, γ and *n* of size 53 *×* 53 and 1 vector *l_x_* of size 53. The equations describe the evolution over time of the level of expression of genes, enzymes as well as concentrations of metabolites and values of reaction flows. **Link to model equations**

**S2 Appendix. Code**. The code is made publicly available at the following link: **Link to Code**

## Acknowledgements

This project has received financial support from the CNRS through the MITI interdisciplinary programs (MetaMod Project 2021-2022). We thank Alaa Tafech for performing the experiments of spheroid growth.

